# Senescent mast cells contribute to the progression of benign prostatic hyperplasia via SCF/c-KIT mediated endothelial-mesenchymal transition

**DOI:** 10.1101/2025.06.02.657200

**Authors:** Tianyu Cao, Jilong Tang, Dapeng Zhou, Chencheng Yao, Junhao Xu, Yi Liu, Chengling Feng, Youwei Shi, Meng Wang, Wenbo Chao, Huanjin Liao, Jin He, Liang Qin, Jiayan Luan, Xiaohai Wang, Di Cui, Yiping Zhu, Yuan Ruan, Shujie Xia, Bangmin Han, Wenhuan Guo, Li Li, Yifeng Jing

**Author notes:** Corresponding author. Correspondences: **Yifeng Jing**, MD, Ph.D. Department of Urology, Shanghai General Hospital, Shanghai Jiaotong University School of Medicine, Shanghai, 200080, China. **Li Li**, MD, Ph.D. Department of Laboratory Medicine, Shanghai General Hospital, Shanghai Jiao Tong University School of Medicine, Shanghai, 200080, China; Department of Laboratory Medicine, Shanghai Clinical Research and Trial Center, ShanghaiTech University, Shanghai, 201203, China. **Wenhuan Guo**, MD, Ph.D. Department of Pathology, Shanghai Ninth People’s Hospital, Shanghai Jiaotong University School of Medicine, Shanghai, 200023, China. These authors have contributed equally to this work (Tianyu Cao, Jilong Tang, Dapeng Zhou).

## Abstract

Benign prostatic hyperplasia (BPH) is a common age-associated urological condition characterized by stromal expansion, but its cellular origins and regulatory mechanisms remain unclear. In this study, we identified endothelial-to-mesenchymal transition (EndMT) as a contributor to stromal cell accumulation in BPH. Using single-cell transcriptomic analysis, endothelial lineage tracing in mice, and validation in human samples, we showed that senescence-associated mast cells increased the expression of vascular endothelial growth factor A (VEGFA) and transforming growth factor-beta 1 (TGF-β1) through the stem cell factor (SCF)/c-KIT–MAPK–JUND signaling pathway, thereby inducing EndMT in endothelial progenitor cells. Stromal fibroblasts express SCF, promoting mast cell activation and establishing a feedback loop that supports continued stromal proliferation. Intriguingly, inhibition of mast cell activation reduces EndMT and attenuates prostate enlargement in vivo. Thus, these findings revealed a senescence-linked immune–stromal interaction in the aging prostate and identify potential targets for therapeutic intervention in BPH.

## Introduction

Benign prostatic hyperplasia (BPH) is one of the most prevalent urogenital diseases in elderly men [1]. Approximately 50% of BPH patients experience moderate to severe lower urinary tract symptoms, which may further lead to serious complications such as renal failure [1]. Although aging is a critical risk factor for the development of BPH, not all elderly men develop to this condition. Clinical observations indicate that some elderly men exhibit only mild prostate enlargement (volume <30 ml), referred to as “small prostate,” while others show significant enlargement (volume >80 ml), termed “large prostate” [2]. Exploring the mechanisms underlying BPH from the perspective of prostate volume differences could provide novel insights and valuable information for pathophysiological research on this disease.

Histopathologically, BPH is characterized by the co-proliferation of glandular epithelium and stroma. Our prior studies revealed that stromal hyperplasia predominates in large prostates, whereas epithelial hyperplasia is more common in small prostates [3]. However, the key factors driving stromal hyperplasia in large prostates remain unclear. Studies have shown that despite significant stromal hyperplasia in large prostates, the proliferative activity of stromal cells was comparable with that in small prostates [4,5], suggesting that the origin of the hyperplastic stromal cells warrants further investigation. It was proposed that epithelial-mesenchymal transition (EMT) contributes to BPH progression, however, this hypothesis remains controversial due to the observation that hyperplastic nodules in many BPH tissues mainly consist of stromal components with minimal epithelial involvement [4].Angiogenesis represents another prominent histological feature of BPH [6]. An alternative hypothesis posits that hyperplastic vascular endothelial cells in BPH tissues could undergo transformation into stromal cells. Early studies reported the loss of vascular endothelial cells in BPH lesions, which implies the potential occurrence of endothelial-mesenchymal transition (EndMT) in these tissues [4]. Notably, EndMT, as a specialized form of cellular transdifferentiation, has been recognized to play a pivotal role in the pathophysiology of various chronic fibrotic diseases [7–9]. Nevertheless, the existence and functional significance of EndMT in BPH remain to be elucidated.

In recent years, the remodeling of the local immune microenvironment in BPH tissues has garnered increasing attention. Aging is associated with dynamic changes in immune cell composition and sustained release of inflammatory factors. Some studies have even proposed that BPH may exhibit autoimmune-like features [10,11]. Among various immune cells, mast cells, as long-lived innate immune cells capable of repeated activation and continuous secretion of inflammatory mediators, remain underexplored regarding their role in BPH pathogenesis. Existing evidence indicates a significant increase in mast cell infiltration in BPH tissues [12], where the IL-6 they secrete activates the STAT3/Cyclin D1 pathway, thereby promoting prostate epithelial proliferation [13]. Furthermore, mast cells have been shown to induce EMT in target cells by secreting factors such as IL-8 and TGF-β in other disease models. For example, Visciano et al. demonstrated that mast cells induce EMT and stemness in thyroid cancer cells via IL-8 [14], while Yin et al. reported that mast cell-derived exosomes mediate EMT in lung epithelial cells [15]. These findings provide critical insights into the potential involvement of mast cells in EndMT in BPH. However, no systematic research has yet elucidated whether mast cells contribute to EndMT in BPH tissues, whether they can induce the transdifferentiation of vascular endothelial cells into mesenchymal cells, or whether their effects are mediated through aging-associated mechanisms. Addressing these key scientific questions is essential for advancing our understanding of BPH pathogenesis.

Here, we investigate the hypothesis that senescent mast cells drive the progression of BPH by inducing EndMT in endothelial progenitor cells. Using single-cell RNA sequencing, lineage tracing in murine models, and functional validation in vitro and in vivo, we show that mast cells in large prostates exhibit features of cellular senescence and secrete VEGFA and TGF-β1, two potent EndMT inducers. These cytokines trigger the transdifferentiation of endothelial cells into myofibroblast-like stromal cells, while stromal cells, in turn, secrete stem cell factor (SCF), amplifying mast cell activation via the SCF–c-KIT–MAPK–JUND axis. Our findings identify a previously unrecognized senescence-driven immune–stromal feedback loop that sustains stromal expansion in the aging prostate, providing a mechanistic explanation for the differential progression of BPH and unveiling potential therapeutic targets.

## Results

### A single-cell atlas of prostate reveals unique cellular in large prostates

To investigate age-related changes and mechanisms underlying BPH development, we conducted single-cell RNA sequencing (scRNA-seq) on transition zone tissues from 6 elderly men with large prostates and 5 elderly men with BPH (aged 60-75 years), excluding those with prostate cancer, using the 10x Genomics Chromium platform. We also obtained scRNA-seq data from 3 young donors from public databases (GSE172357) [16] (Fig.1A). After quality control, we processed a total of 104,547 cells. Unsupervised clustering identified and visualized eight major cell types: epithelial, endothelial, fibroblasts, T cells, B cells, NK cells, mast cells, and myeloid cells (Fig.1B, S1). Subgroups within these cell types were further annotated (Fig.1C, S2). Characteristic genes, expression levels, and detected genes/UMIs for each subgroups were visualized using heatmaps, UMAP plots, and violin plots (Fig.S3-6). We observed significant differences in cellular composition among normal, large, and small prostates. Large prostates exhibited higher numbers of fibroblasts and endothelial cells compared to small and normal prostates; normal prostates contained the highest proportion of epithelial cells. Elderly patient prostates were enriched with T cells, NK cells, B cells, and myeloid cells relative to normal prostates (Fig.1D, S7). These findings were validated using an independent cohort of 20 large, 20 small, and 12 normal prostate tissues (Fig.1E, S8A). We performed H&E staining and statistical analysis on 50 large prostate samples and 46 small prostate samples, revealing a significantly higher proportion of stromal components in the transition zone of large prostates compared to small prostates. (Fig.1F).

**Figure 1.**
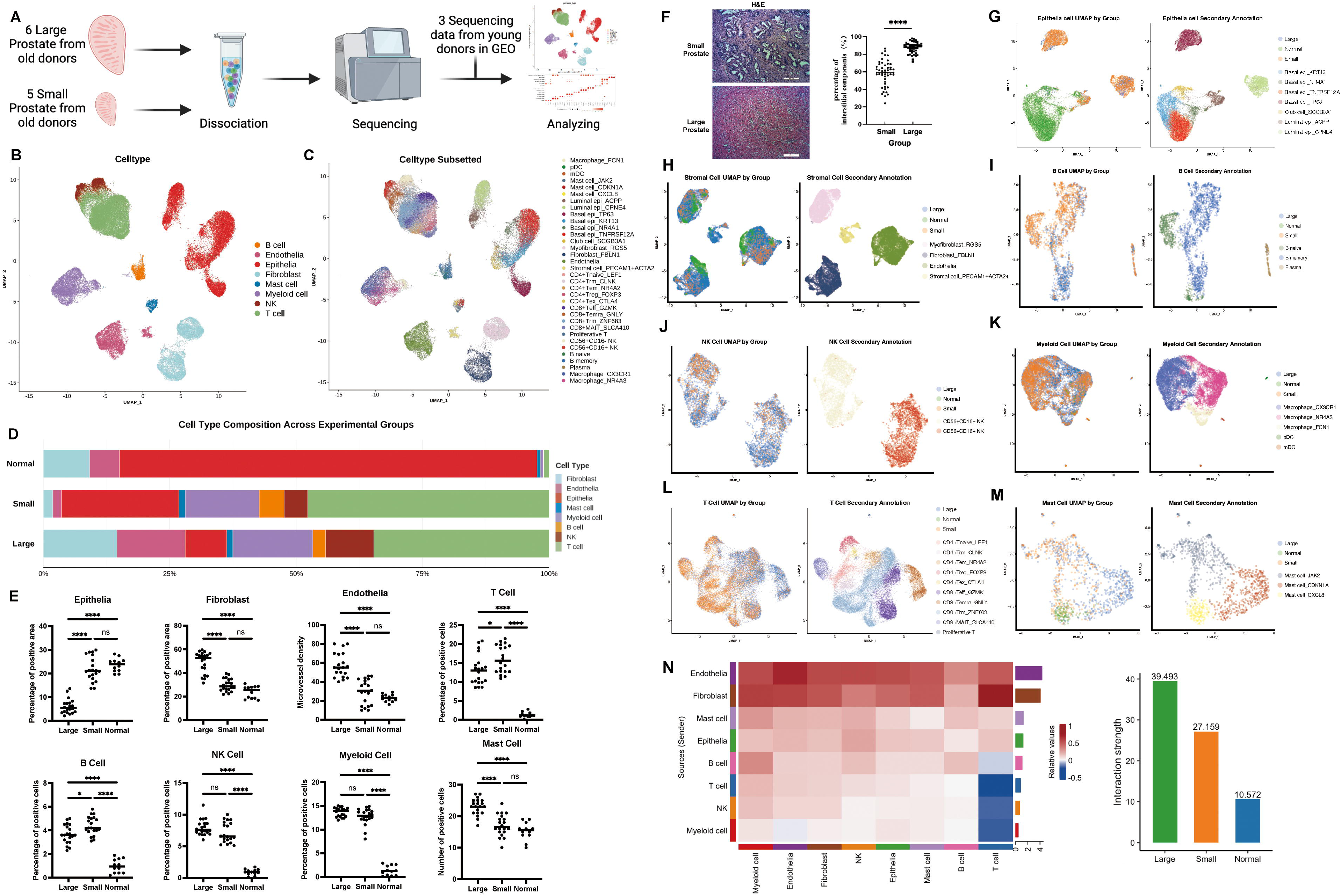
Single-cell transcriptomic atlas reveals cellular heterogeneity and aging phenotypes in large human prostates. (A) Workflow of scRNA-seq across normal, small, and large prostates. (B-C) UMAP plot of annotated cell types in the human prostate. (Color-coding corresponds to distinct cell type). (D) Stacked bar plot quantifying the relative abundance of cell clusters within each experimental group. (E) Scatter plot correlating cluster proportions with experimental group stratifications. (Large and small, n=30; Normal, n=12) (F) Histological examination images of large and small prostate tissue stained with H&E, along with statistical graphs of the stromal components of the prostate. (Each group, n=60) (G-M) UMAP plot of each cell subsets with cluster-specific or group specific color annotation. (N) Bar plot summarizing interaction frequencies between major cell clusters across experimental groups, and heatmap showing differential number of interactions among cell subsets across three experimental groups. *mDC□=□myeloid dendritic cell; MAIT□=□mucosal-associated invariant T cell; NK□=□natural killer; pDC□=□plasmacytoid dendritic cell; Tem□=□Effector memory T cell; Treg□=Regulatory T cell, Trm□=□T resident memory T cell ; Temra=Terminally differentiated effector memory T cell; UMAP, uniform manifold approximation and projection; H&E, hematoxylin and eosin; ns p* ≥ *0.05, * p < 0.05, *** p < 0.001, **** p < 0.0001*.

We initially focused on analyzing the non-immune cell subpopulations within the prostate tissue. Epithelial cells were categorized into two major types: luminal and basal cells, with a total of seven subgroups identified. Our results showed that *CPEN4*+ luminal cells and *TP63*+ basal cells were specifically enriched in elderly prostates, whereas *KRT13*+/*NR4A1*+/*TNFRSF12A*+ basal cells were predominantly enriched in normal prostates. Additionally, club cells and *ACPP*+ luminal cells were commonly enriched in both elderly and normal prostates (Fig.1G, S8B). Notably, no specific epithelial cell subgroups were observed between large and small prostates, suggesting that age is the primary factor driving epithelial cell changes rather than BPH progression. We classified the stromal components in the prostate into subgroups and found that stromal cells were predominantly enriched in large prostates (Fig.1H, S8B). However, cell cycle and Ki67 analysis revealed no significant differences in the proportions of G2M phase fibroblasts and epithelial cells between large and small prostates, although the proportion of G2M phase endothelial cells was notably higher in large prostates (Fig.S9A-D). This suggests that in larger prostates, the cells exhibiting a higher proliferative potential are endothelial cells rather than fibroblasts. Notably, we identified a novel *PECAM1*+*ACTA2*+ stromal cell subgroup in large prostates, which was nearly absent in small prostates (Fig. 1H). This cell population concurrently displays characteristics of both endothelial cells and fibroblasts, suggesting a distinct and unique cellular identity. We further validated this finding using a public dataset of BPH (GSE145928 and GSE183676) (Fig. S9E-F) [17,18].

Aging facilitates the infiltration of immune cells from peripheral sites into tissues [19]. We performed a clustering analysis of immune cells specifically identified within the prostate tissue . NK cells and B cells showed quantitative differences but not in subgroup composition (Fig.1I-J). Myeloid cells did not exhibit significant differences in either aspect (Fig.1K). Consistent with our previous results [3], effector T cell subsets expressing granzyme (CD8+Teff_GZMK and CD8+Temra_GNLY) were significantly enriched in larged prostates (Fig.1L). Differential gene enrichment analysis revealed upregulation of pathways related to cell adhesion, antigen presentation, and cytotoxicity in T cells from large prostates (Fig.S10A-B). Mast cells are evolutionarily conserved, long-lived innate immune cells, and consistent with previous studies [20,21], our clustering analysis revealed significant heterogeneity among mast cell subgroups, with distinct compositions observed in large, small, and normal prostates (Fig. 1M). Specifically, a CDKN1A-expressing mast cell subgroup was enriched in large prostates. This suggests that prostates of larger size possess a distinctively unique composition of the immune microenvironment.

Using CellChat, we analyzed intercellular communication and found that large prostates exhibited the highest intensity and number of intercellular communications, followed by small prostates, with normal prostates showing the least (Fig.1N, S10C). This aligns with the continuous hyperplasia in large prostates and the altered microenvironment in elderly prostates. Endothelial cells, fibroblasts, and mast cells were the most prominent cell types with enhanced intercellular communication in large prostates. Significantly upregulated pathways included androgen-related (e.g., testosterone, DHT), immune enrichment-related (e.g., ICAM, VCAM, CXCL, CCL), and secretory factor-related (e.g., VEGF, EGF, TGFβ, KIT) pathways (Fig.S10D). These results highlight the complex microenvironment in elderly and large prostates.

### The large prostate displays distinct cellular senescence-associated phenotypic features

We used the aging gene set (SAUL_SEN_MAYO) to assess the overall aging status of normal, large and small prostates [22]. The results showed that only large prostates exhibited significant aging phenotypes (Fig.2A). We identified senescence-related differentially expressed genes (DEGs) in aged prostates, revealing pronounced enrichment of senescence genes in epithelial, stromal, and mast cells (Fig.2B, S11A). This is consistent with the results of the prior intercellular communication analysis, revealing marked alterations in epithelial cells, stromal cells, and mast cells within the large prostate. CDKN1A is one of the most commonly used markers for cellular senescence [23], and CDKN1A staining confirmed a higher number of CDKN1A-positive cells in large prostates (Fig.2C-D), highlighting their characteristic senescence phenotype.

**Figure 2.**
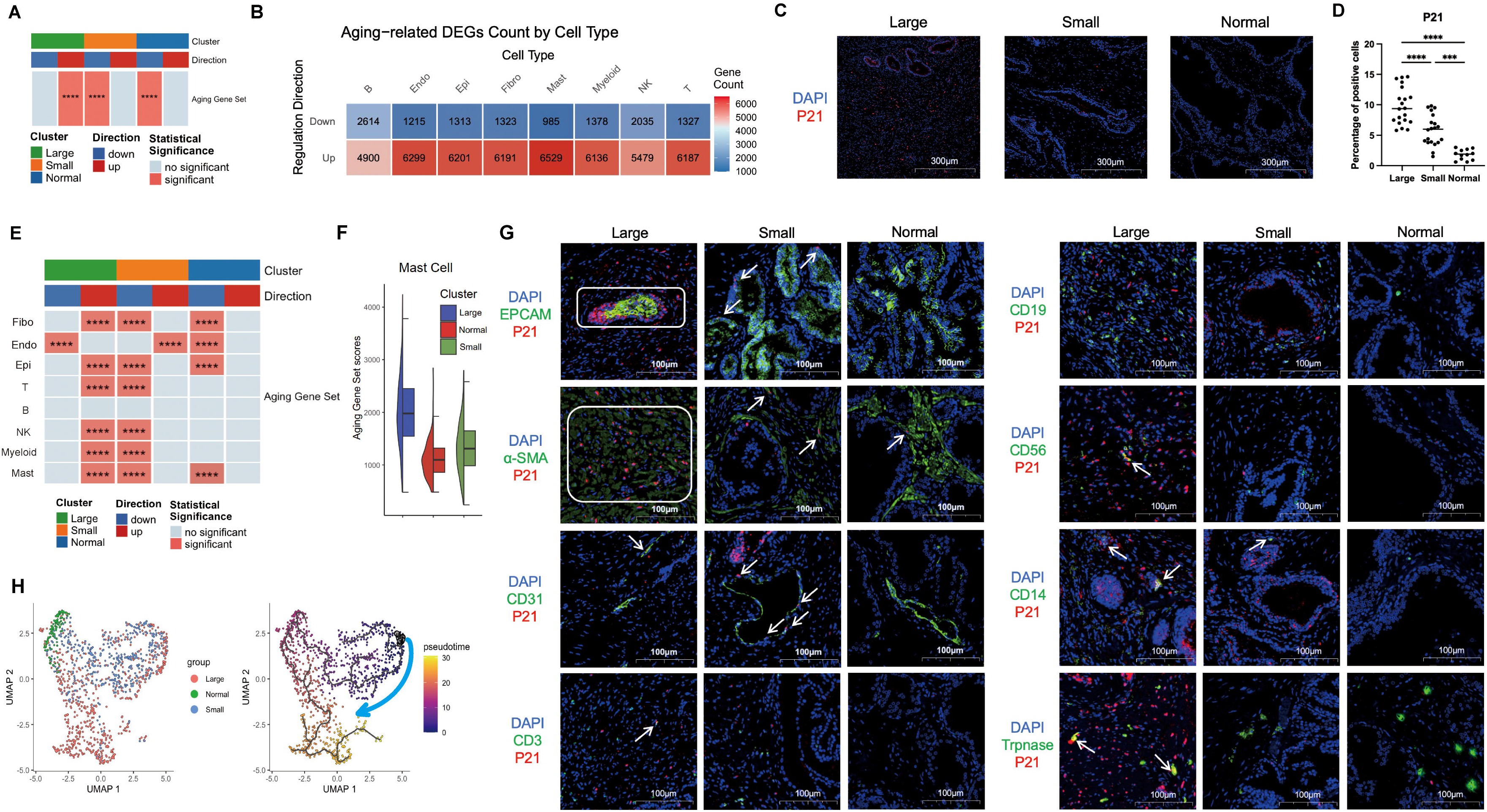
Cellular aging-related phenotypic features of the large prostate. (A) Comparative aging scores across experimental groups calculated using the SAUL_SEN_MAYO senescence gene set (GSEA ID: M45803). (B) Heatmap of differentially expressed genes (DEGs) identified in all cell subsets through scRNA-seq analysis. (C-D) Immunofluorescence staining shows the expression of P21 in different experimental groups of human prostate (Large and small, n=30; Normal, n=12). (E) The senescence score for each cell was calculated using the SAUL_SEN_MAYO senescence gene set, providing quantitative insights into cellular senescence. (F) The bar plot presents the mast cell senescence score results calculated based on the SAUL_SEN_MAYO aging gene set. (G) Multicolor immunofluorescence staining was employed to visualize the expression levels of P21 in epithelial cells (EPCAM), endothelial cells (CD31), stromal cells (α-SMA), T cells (CD3), B cells (CD19), NK cells (CD56), myloid cells (CD14) and mast cells (Trpnase) within the human prostate. (H) Pseudo-time trajectory analysis reveals dynamic differentiation states of mast cells. *\*\*\* p < 0.001, **** p < 0.0001*.

We further employed single-cell data and human prostate tissue sections to systematically analyze the senescence characteristics of various cell types in the prostate. In large prostates, only endothelial cells maintained a non-senescent phenotype, whereas fibroblasts, epithelial cells, T cells, B cells, NK cells, myeloid cells, and mast cells exhibited more pronounced senescent phenotypes (Fig.2E-G). Small prostates and those from young individuals displayed similar senescence patterns. Notably, we found that mast cells were the sole immune cell type in large prostates that not only exhibited senescence but also increased in number. Moreover, mast cells were the only immune cells enriched in both young and old prostates. We observed that mast cells in large prostates specifically upregulated cytokines mRNA Expression such as *VEGFA*, *NAMPT* and *TGFB1* (Fig.S11B). Through pseudo-time trajectory analysis, we further confirmed that mast cells in large prostates were in a terminal differentiation state, while mast cells in normal prostates remained at an intermediate stage and lacked the expression of secreted cytokines like VEGFA (Fig.2H, S11C-D). Based on these findings, we speculate that mast cell senescence may play a critical role in the progression of benign prostatic hyperplasia.

### Mast cells drive endothelial progenitor cells to undergo EndMT and exacerbate the progression of BPH

We previously identified a unique cell population, *PECAM1*+*ACTA2*+ stromal cell, that is exclusively present in large prostates exhibiting senescence-associated phenotypes. Furthermore, we confirmed that CD31 (encoded by *PECAM1*)+ α-SMA (encoded by *ACTA2*)+ stromal cells are specifically localized to larger prostate tissues (Figure 3A). Under pathological conditions, endothelial cells undergo remodeling, gradually losing specific protein expression and acquiring characteristics of stromal cells, including morphology and function—a process known as EndMT [24]. EndMT plays a critical role in various pathophysiological processes of fibrotic diseases, leading to myofibroblast activation, increased production of pro-fibrotic molecules, enhanced tissue stiffness, and progression of fibrosis [25]. Previous studies have noted thickened vascular walls and loss of endothelial layers in BPH stromal tissues, suggesting the occurrence of EndMT, though this has not been fully confirmed [4]. Based on these observations, we hypothesized that EndMT occurs during BPH progression and confirmed through pseudo-time trajectory analysis that *PECAM1*+*ACTA2*+ stromal cells originate from endothelial cells (Fig.3B, S12A).

**Figure 3.**
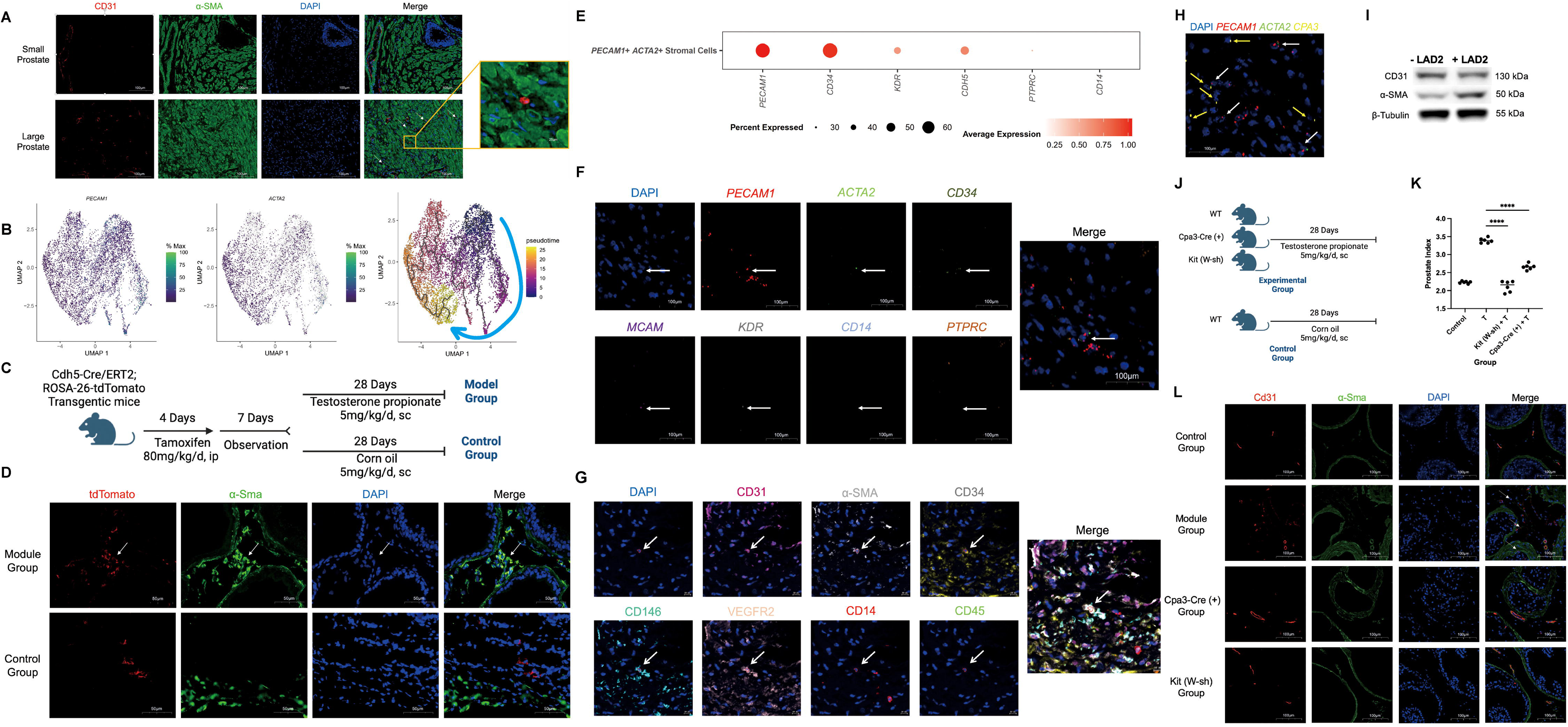
Senescent mast cells trigger EndMT and exacerbate the progression of BPH. (A) Multicolor immunohistochemical staining of the transitional zone tissue of the prostate reveals the presence of CD31+ α-SMA+ cells within the large prostate. (B) Pseudotime trajectories of endothelia cell reveal *PECAM1+ ACTA2+* stromal cell originate from endothelia cells. (C) Schematic diagram of the experimental steps of endothelial cell tracing mice. (D) Cd31+ tdTomato+ cells exist in BPH tissues of testosterone propionate-induced, endothelial cell-tracing mice. (E) The dot plot illustrates the expression profile of ECFC marker genes within the *PECAM1+ ACTA2+* cells population in single-cell RNA sequencing. (F) RNAScope has confirmed that the *PECAM1+ ACTA2+* cells exhibit the *CD34+, MCAM+, KDR+, CD14- and PTPRC-* phenotype. (G) TissueFAXS Spectra presence of CD31+ α-SMA+ CD34+ CD146+ VEGFR2+ CD14-CD45-cells in large prostates. (H) RNA-Scope reveals the spatial relationship between mast cells (*CPA3*+ cells) and PECAM1+ ACTA2+ cells. (I) Western-blot showed that LAD2 cells could promote ECFCs to undergo EndMT. (J) Schematic diagram of the experimental steps of mast cell-deficient mice. (K) Prostate index of control group mice, model group mice, and mast cell-deficient mice. T represents daily subcutaneous injection of 5 mg/kg/d testosterone propionate. (L) The multicolor immunohistochemical staining of mouse prostate tissue reveals the presence of Cd31+ α-Sma+ cells. *\*\*\*\* p < 0.0001*.

We further validated through vivo experiments whether CD31+α-SMA+ stromal cells were generated via EndMT of endothelial cells. Testosterone propionate-induced BPH is a commonly used animal model that leads to the proliferation of both epithelial and stromal cells in the prostate [26]. In testosterone propionate-induced BPH mice, we observed Cd31+α-Sma+ stromal cells (Fig.S12B), and high-dose testosterone induced cellular senescence in the prostate (Fig.S12C). To track endothelial cells, we generated Cdh5-Cre/ERT2;ROSA26-tdTomato transgenic mice (Fig.3C). After tamoxifen administration, all endothelial-derived cells expressed tdTomato. Compared with the control group, the BPH model mice exhibited a significant presence of tdTomato+α-Sma+ stromal cells (Fig.3D), confirming that CD31+α-SMA+ stromal cells arise through EndMT and are critically involved in the progression of BPH.

Given the growth and differentiation potential of endothelial cells in large prostates, we inferred that endothelial cells undergoing EndMT exhibit an endothelial progenitor cell phenotype. Most endothelial cells in the prostate express endothelial colony-forming cells (ECFCs) phenotype (Fig.3E, S12D), suggesting that these cells are primarily ECFCs rather than myeloid angiogenic cells. ECFCs can form new blood vessels in vivo and contribute to vascular repair [27–29]. This finding was also validated in published datasets of BPH (GSE145928 and GSE183676) (Fig.S12E-F). Using RNAscope™ HiPlex, we detected *PECAM1*+*ACTA2*+*CD34*+*MCAM*+ *KDR*+*CD14*-*PTPRC*-cells in large prostates (Fig.3F). The presence of CD31+α-SMA+CD34+CD146+VEGFR2+CD14-CD45-cells in large prostates was further confirmed using TissueFAXS Spectra (Fig.3G), supporting the occurrence of EndMT within BPH-associated ECFCs.

Prior research has demonstrated that mast cells are spatially located near blood vessels and secrete TGF-β1, facilitating EMT in respiratory epithelial cells [15,30]. Using RNAscope™ HiPlex and mIHC, we demonstrated that mast cells in BPH tissues were closely associated with *PECAM1*+*ACTA2*+ stromal cells in human large prostate (Fig. 3H, S13A-B). We previously demonstrated that mast cells in large prostates exhibit elevated expression levels of *VEGFA* and *TGFB1*. VEGFA is a critical cytokine for neovascularization and has emerged as an important target for anti-angiogenic therapy [24], while TGF-β1 is the most potent inducer of EndMT known to date [7,24]. These findings suggest that mast cells may promote EndMT through paracrine signaling mechanisms. To test this hypothesis, we treated ECFCs, derived from peripheral blood CD34+ cells, with conditioned medium from the human mast cell line LAD2. The results revealed that LAD2-conditioned medium significantly reduced CD31 expression in ECFCs while markedly enhancing α-SMA expression (Fig.3I), thereby confirming the ability of mast cells to induce EndMT in ECFCs. We futher generated two mast cell-deficient transgenic mouse strains: Kit (W-sh) and Cpa3-Cre (+) (Fig.S13C) [31,32]. After inducing BPH (Fig.3J), wild-type mice developed prostatic hyperplasia characterized by the presence of Cd31+α-Sma+ stromal cells (Fig.3K-L), whereas mast cell-deficient mice exhibited significantly smaller prostates without these stromal cells. Additionally, mast cell-deficient mice showed reduced P21 expression (Fig.S13D), lower microvascular density, and decreased immune cell infiltration in the prostate (Fig.S13E). Collectively, these findings provide robust and compelling evidence that mast cells are critically involved in driving the progression of BPH and EndMT within BPH pathophysiology.

### SCF/c-KIT pathway activates mast cells and mediates EndMT in prostate

Mast cells primarily possess two types of receptors: c-KIT and FcεRI [31,33]. Using Fcer1a-KO mice, we demonstrated that the activation of mast cells in BPH does not depend on the canonical IgE-FcεRI-related ITAM signaling pathway (Fig.S14A). Consequently, we focused our investigation on the c-KIT pathway. The c-KIT pathway in prostate is primarily composed of the ligand SCF and the receptor c-KIT (Fig. 4A). SCF, a dimeric molecule existing in both membrane-bound and soluble forms, is predominantly expressed by fibroblasts and endothelial cells throughout the body and plays a critical role in the survival, development, and activation of mast cells. Notably, we observed significantly higher expression levels of SCF in large prostates compared to small prostates (Fig.4B). This finding suggests that SCF secreted by prostate fibroblasts may serve as a key factor in activating mast cells.

**Figure 4.**
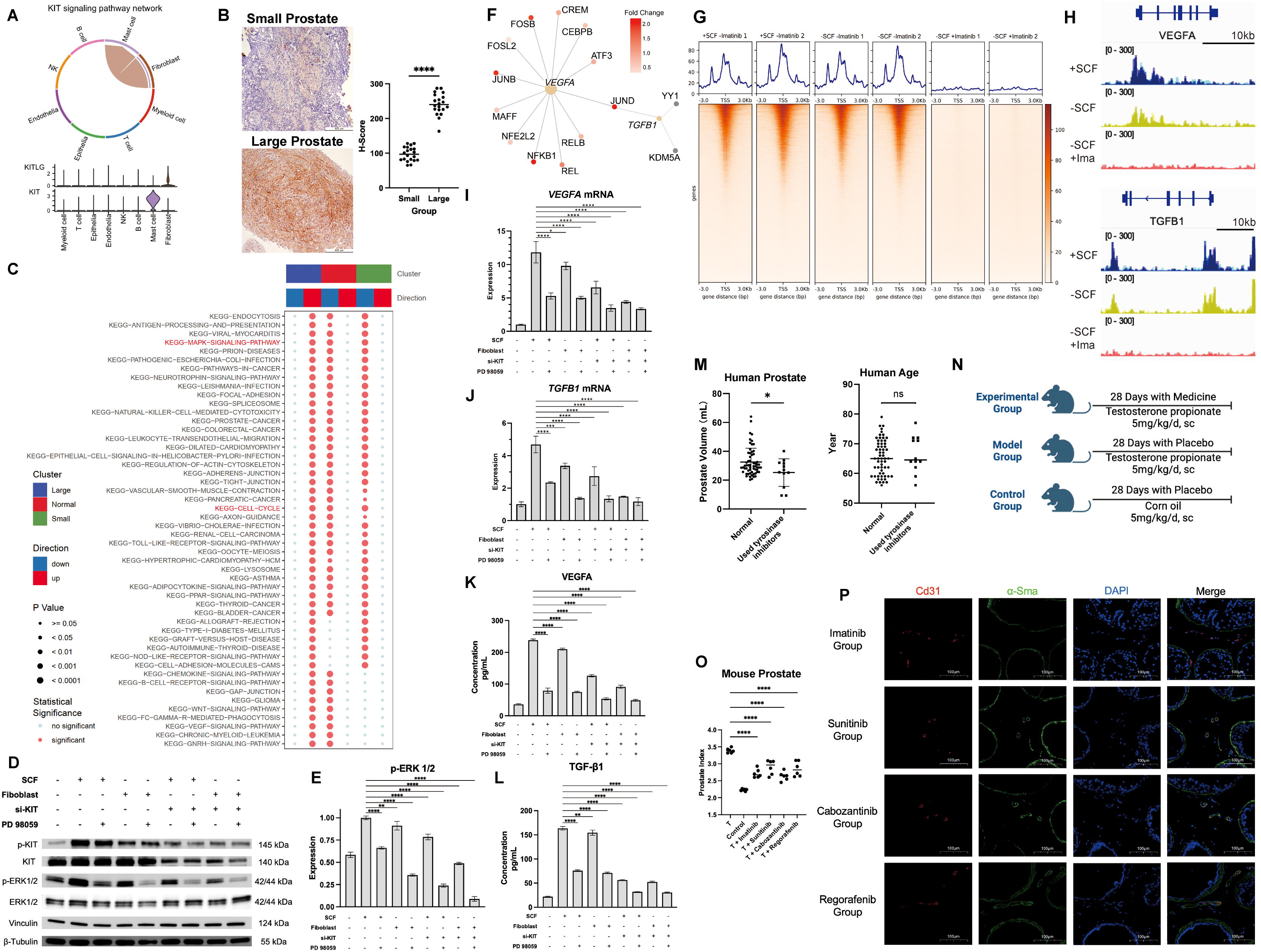
SCF/c-KIT pathway activates mast cells and mediates EndMT in prostate. (A) CellChat analysis revealed that stromal cells are the main SCF secreting cells. (B) Immunohistochemical staining was performed on large and small prostate tissues using anti-SCF. (Each group, n=20) (C) Functional enrichment heatmap of Mast cell DEGs across experimental groups, analyzed by irGSEA (ssGSEA method, FDR < 0.05). (D-E) Western-blot revealed the effects of fibroblasts, SCF, and PD98059 on the KIT and MAPK pathways in mast cells. (F) The dot plot shows the transcription factors JUND that regulate *VEGFA* and *TGFB1* in mast cells in prostate tissue. (G-H) The CUT&Tag analysis of mast cells confirmed that SCF could promote *VEGFA* and *TGFB1* transcription through JUND in the living mast cells. (I-J) qPCR revealed the effects of fibroblasts, SCF, and PD98059 on the expression of *VEGFA* and *TGFB1* mRNA in mast cells. (K-L) Elisa reveals the effects of fibroblasts, SCF, and PD98059 on the secretion of VEGFA and TGF-β1 by mast cells. (M) Comparison of prostate volume and age between elderly men taking TKIs and normal elderly men. The data on prostate volume and age of normal elderly men were obtained from the physical examination center. (N) Schematic diagram of the experimental steps of mice. (O) Prostate index of mice in the control group, model group, and TKIs-treated group. T represents daily subcutaneous injection of 5 mg/kg/d testosterone propionate. (P) The multicolor immunohistochemical staining of mouse prostate tissue reveals the presence of Cd31+ α-Sma+ cells. *ns p* ≥ *0.05, * p < 0.05, ** p < 0.01,*** p < 0.001, **** p < 0.0001*.

Through gene enrichment analysis, we identified the MAPK pathway as the most significantly upregulated pathway in large prostates (Fig.4C), a result confirmed by Gene Set Enrichment Analysis (GSEA) (Fig.S14B). To further validate this observation, we cultured primary stromal fibroblasts expressing SCF from large prostates (Fig.S14C) and co-cultured them with LAD2 cells, using SCF-treated cells as a positive control. Consistent with scRNA-seq data, we found that SCF secretion by stromal fibroblasts from large prostates activated c-KIT on the surface of LAD2 cells, thereby promoting MAPK pathway transduction within LAD2 cells. Interfering with c-KIT expression in LAD2 cells using siRNA or treating with the MEK inhibitor PD 98059 effectively inhibited MAPK pathway transduction (Fig.4D-E, S14D).

Integrating findings from previous experiments, we confirmed that mast cells promote EndMT. SCENIC transcription factor analysis revealed that the expression of VEGFA and TGFB1 is jointly regulated by the transcription factor JUND (Fig.4F). As a key component of the activator protein-1 (AP-1) complex, JUND is modulated by multiple signaling pathways, including MAPK and PI3K [33]. CUT&TAG analysis further corroborated that in mast cells, the c-KIT pathway enhances the transcription of VEGFA and TGFB1 via JUND (Fig.4G-H). Adding SCF to the LAD2 culture medium or co-culturing LAD2 cells with stromal fibroblasts from large prostates increased the transcription of VEGFA and TGFB1 mRNA in LAD2 cells, ultimately leading to the secretion of VEGFA and TGF-β1. Both effects were inhibited by interfering with c-KIT expression in LAD2 cells using siRNA or treating with PD 98059 (Fig.4I-L).

Tyrosine kinase inhibitors (TKIs), which inhibit c-KIT-related signaling pathways [34], have prompted speculation regarding their potential application in BPH treatment. TKIs are also first-line drugs for chronic myeloid leukemia (CML) [35]. We observed that stable elderly male CML patients exhibited reduced prostate volume after taking TKIs compared to the normal population (Fig.4M), suggesting that TKIs can suppress BPH progression. In animal experiments, we confirmed that four commonly used clinical TKIs—imatinib, sunitinib, cabozantinib, and regorafenib—effectively slowed BPH progression (Fig.4N-O), reduced the number of Cd31+α-Sma+ stromal cells in the prostate (Fig.4P), while inhibiting angiogenesis, reducing the infiltration of Cd45+ immune cells, and alleviating prostate aging (Fig.S14E-F). However, it is important to note that while TKIs demonstrate therapeutic efficacy, they may cause side effects such as rash and diarrhea [36]. Although no deaths occurred in TKI-treated mice, differences in weight loss were observed compared to the control group (Fig.S14G). Therefore, we conclude that the use of TKIs for treating BPH, a benign condition, carries potential risks.

### Mast cell membrane stabilizers and TGF-**β** signaling pathway inhibitors suppress EndMT in prostate and BPH progression

In light of the significant side effects associated with TKIs and their potential risks in clinical application [36], we have identified four mast cell membrane stabilizers and TGF-β signaling pathway inhibitors with fewer side effects as potential candidates for BPH suppression. Tranilast, a mast cell membrane stabilizer and inhibitor of the TGF-β signaling pathway, is frequently employed to treat allergic conditions, with some researchers exploring its use for fibrotic diseases [37,38]. Cromolyn sodium, a classic mast cell membrane stabilizer, is among the most frequently used medications in the clinical treatment of allergic diseases, also shows certain application prospects in fibrotic diseases [39,40]. Ketotifen, a non-competitive H1 antihistamine and mast cell membrane stabilizer, typically prescribed for allergic conditions, is now being researched for its potential use in treating conditions such as irritable bowel syndrome and tumors, which may be associated with mast cells [41–43]. Asiaticoside, an inhibitor of the TGF-β signaling pathway, possesses anti-fibrotic and anti-inflammatory properties and is clinically used to treat fibrosis-related diseases and scars [44,45]. Due to their proven clinical safety in applications, we decided to further explore their potential uses in BPH suppression.

Initially, we tested the inhibitory effects of these four drugs on EndMT in *vitro*. We incorporated the four drugs into a novel mast cell culture medium and, after culturing the mast cells for 48 hours, transferred the supernatant into ECFC culture medium. We observed that these drugs suppressed EndMT in *vitro* (Fig.5A). Subsequently, we confirmed this finding in *vivo*. We observed that these four drugs effectively reduced testosterone propionate-induced BPH in mice (Fig.5B-C), decreased the number of Cd31+ α-Sma+ stromal cells in the prostate (Fig.5D),, while inhibiting angiogenesis, reducing the infiltration of Cd45+ immune cells, and alleviating prostate aging (Fig.S15A-B). Notably, tranilast has exhibited efficacy comparable to that of finasteride (Fig.S15C).None of the four drugs caused mortality in mice, and we noted that, with the exception of ketotifen and cromolyn sodium, the administration of the other two drugs (tranilast and asiaticoside) did not significantly decrease mouse body weight (Fig.S15D).

**Figure 5.**
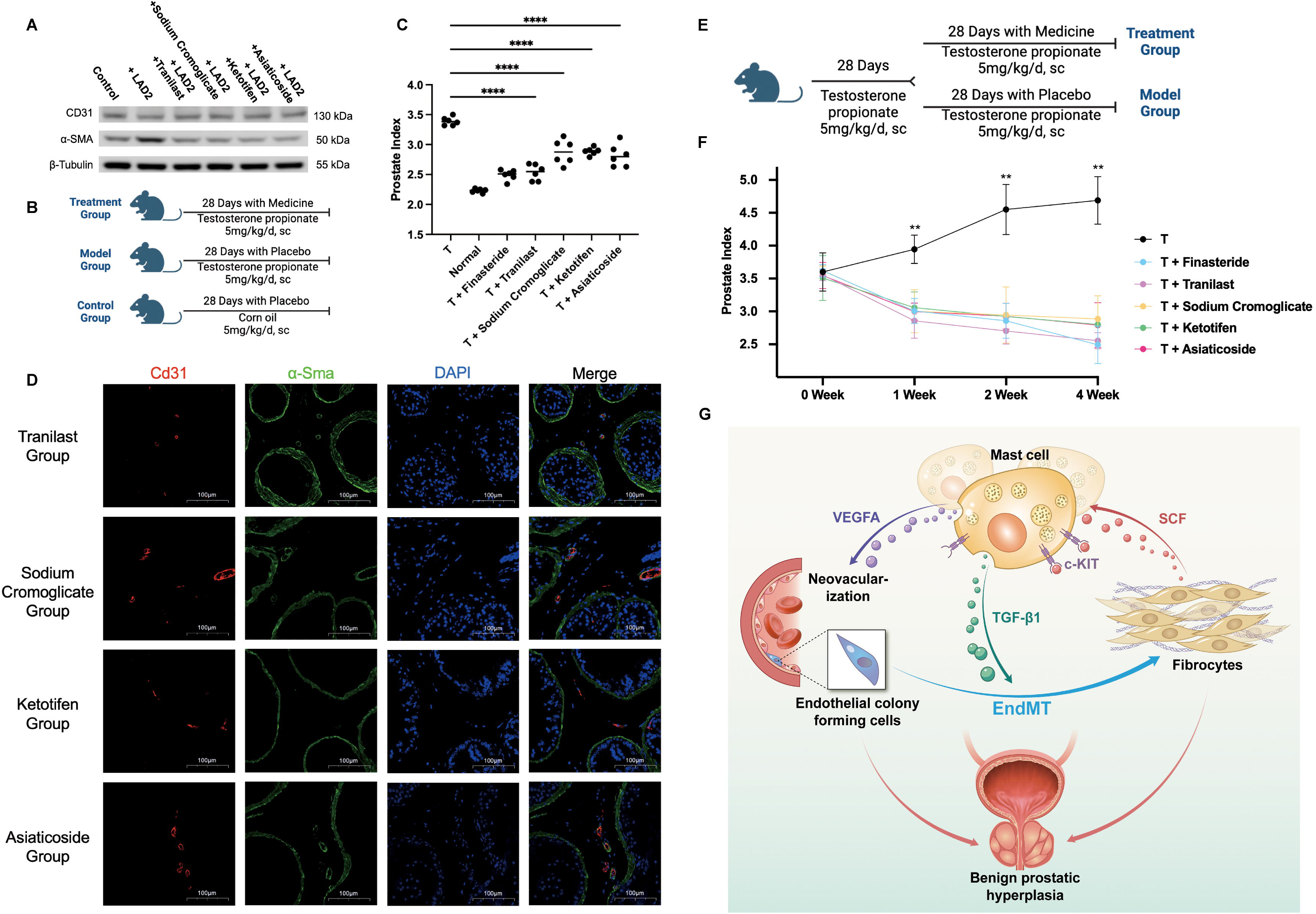
Mast cell membrane stabilizers and TGF-**β** signaling pathway inhibitors suppress EndMT in prostate and BPH progression. (A) Mast cell membrane stabilizers and TGF-β inhibitors can inhibit the promotion of EndMT in ECFC by LAD2 cells. (B) Schematic diagram of the experimental steps of prevention group mice. (C) Prostate index of mice in control group, model group, mast cell membrane stabilizer and TGF-β inhibitor prevention group. T stands for daily subcutaneous injection of 5mg/kg/d testosterone propionate. (D) The multicolor immunohistochemical staining of mouse prostate tissue reveals the presence of Cd31+ α-Sma+ cells. (E) Schematic diagram of the experimental steps of treatment group mice. (F) Prostate index of mice in different treatment group. T stands for daily subcutaneous injection of 5mg/kg/d testosterone propionate. (G) Graphical abstract. *\*\* p < 0.01, **** p < 0.0001*.

To further determine whether the aforementioned four drugs have therapeutic effects on BPH, we administered these four drugs to BPH mice that had been successfully modeled with testosterone propionate, and continued to administer testosterone propionate during the administration period (Fig.5E). The results indicated that all four drugs could continuously inhibit the increase in prostate volume induced by testosterone propionate(Fig.5F). Notably, tranilast had an efficacy comparable to that of finasteride without significantly reducing the body weight of mice (Fig.S15E). In summary, the treatment of BPH with mast cell membrane stabilizers and inhibitors of the TGF-β signaling pathway can inhibit its progression. In comparison to TKIs, these treatments exhibit fewer side effects and are thus considered safer for the organism.

## Discussion

Single-cell sequencing technology has revolutionized our understanding of diseases with unprecedented resolution. In this study, inspired by the clinical observation that prostate volume progression varies significantly among elderly individuals, we performed scRNA-seq on the transition zone of the prostate in elderly patients without prostate cancer, focusing on samples with different prostate volumes. By integrating these data with publicly available datasets, we constructed unbiased atlases of the prostate across different ages and volumes. This not only deepens our understanding of the biological mechanisms underlying BPH progression but also provides a valuable resource for future BPH research.

Previous studies have demonstrated that senescence occurs in the glandular epithelium of enlarged prostates. Meanwhile, earlier studies generally assumed that senescent cells were absent in the prostate stroma[46,47]. Our research reveals that senescence in large prostates extends to epithelial cells, stromal cells, and immune cells, thereby providing critical aging-related insights for future BPH investigations. Furthermore, we were the first to uncover the presence of EndMT in large-volume prostates, which contributes to the formation of CD31+α-SMA+ stromal cells. This finding elucidates the thickening of vascular walls and the loss of the endothelial layer observed in BPH interstitial tissue. CD31+α-SMA+ stromal cells further promote pathological fibrosis, initiate extracellular matrix remodeling, and alter the mechanical properties of the tissue matrix [24], thereby exacerbating BPH-related symptoms. This also explains why some patients exhibit poor responses to drug treatments [48], primarily because existing drugs fail to inhibit the continuous growth of fibroblasts in BPH. We identified ECFCs as potential precursors of EndMT, a phenomenon rarely reported in EndMT studies [49]. ECFCs and their derivatives display immune privilege characteristics, offering protection against homologous antibody/complement-mediated cytotoxicity and T-cell activation-induced lysis [50]. Further investigation is warranted to determine whether this phenomenon is specific to benign BPH or extends to other pathological and physiological processes involving EndMT.

Mast cells are known to induce chronic inflammation and pathological tissue remodeling [51,52]. Our findings reveal that mast cells are the sole immune cells persisting from the mature to the elderly stages of the prostate and serve as the most critical immune cells driving BPH progression. While previous studies have noted an increase in mast cell numbers in BPH [12], the precise role of mast cells in this condition remains underexplored. We demonstrated that elevated expression of SCF in the prostate stroma attracts mast cell progenitors from blood vessels to the prostate stroma, promoting mast cell development, activation, and proliferation [30]. This process represents a key step in angiogenesis and EndMT during BPH progression. Subsequently, EndMT generates new activated fibroblasts, establishing a positive feedback loop between mast cells and fibroblasts, which drives the continuous proliferation of the prostate stroma. Additionally, we revealed that c-KIT activation in mast cells enhances the transcription factor JUND, which upregulates *VEGFA* and *TGFB1* transcription, ultimately leading to the secretion of VEGFA and TGF-β1. This mechanistic insight expands our understanding of mast cells, demonstrating that c-KIT can independently activate mast cells in non-allergic diseases.

Given the incomplete understanding of BPH pathogenesis, the development of drug treatments has been relatively stagnant over the past decades. Currently, 5α-reductase inhibitors and α1-adrenergic blockers remain the primary clinical options [53–55]. Our innovative approach using mast cell membrane stabilizers (cromolyn sodium and ketotifen), TGF-β pathway inhibitor (asiaticoside), and dual-action drug (tranilast) demonstrates their ability to inhibit EndMT in vitro and prevent testosterone-induced BPH in mice. Although these four drugs exhibit less efficacy compared to TKIs in inhibiting BPH in mice, they produce significantly fewer side effects. Using a therapeutic model, we verified the efficacy of these drugs in treating BPH, with tranilast showing comparable effects to the 5α-reductase inhibitor finasteride. Recent reports indicate that sodium tryptophan alleviates lower urinary tract symptoms (LUTS) caused by bacterial infections, while ketotifen reduces fibrosis symptoms induced by mast cells under fibrotic conditions [42,43,52]. These findings suggest that mast cell membrane stabilizers and TGF-β pathway inhibitors, particularly tranilast, hold great promise for BPH treatment, warranting further clinical trials to validate tranilast’s role in managing BPH.

This study is the first to identify mast cell senescence and endothelial-mesenchymal transition (EndMT) in benign prostatic hyperplasia (BPH), revealing the role of mast cells in disease progression and their activation mechanisms. However, several limitations remain. The exact mechanisms underlying BPH initiation remain unclear, particularly the initiating factors and the specific cell populations responsible for activating stromal cells or mast cells. Although this study highlights SCF as a critical factor influencing mast cells, elucidating the differential SCF expression between large and small prostates remains challenging. Future research should focus on determining the precise mechanisms governing increased SCF expression in large prostates.

## Star methods

### scRNA-seq of prostatic cells

After obtaining informed consent from the patient, the Ethics Committee of Shanghai General Hospital approved the collection of human prostate tissue. The large prostate tissue was obtained from male patients undergoing thulium laser enucleation of the prostate (ThuLEP) for BPH, with a preoperative prostate volume exceeding 80 mL under transabdomincal ultrasound. The small prostate tissue was obtained from male patients undergoing ThuLEP for bladder neck contracture, with a preoperative prostate volume measured by transabdominal ultrasound less than 30 mL. All patients did not receive any treatment related to benign prostatic hyperplasia before surgery, and no prostate cancer was found in the postoperative pathology.

Tissues used for scRNA-seq were minced and digested with 1000 U/mL collagenase I + 1 mg/mL DNase I + 1% P/S (Penicillin/Streptomycin) at 37°C for 4 hours, and then treated with the aforementioned TrypLE Express reagent. After cell washing and red blood cell lysis, debris was removed from the cell suspension using a debris removal solution.

According to the manufacturer’s instructions, the single-cell suspension was subjected to generation and preparation of a 10X Chromium chip single-cell library using a 10X Chromium 3’ solution (V2 kit) to capture 2500-12000 cells/channel. Sequencing was performed on the Illumina HiSeq4000 platform (samples L1-L3, S1-S3) and the Illumina HiSeq6500 platform (samples L4-L6, S4-S5). After sequencing, the BCL file was demultiplexed into Fastq files using CASAVA.

### Raw data processing and quality control

The Cell Ranger software pipeline, offered by 10×Genomics, is utilized to demultiplex cell barcodes, align reads to the genome and transcriptome using the STAR aligner, and, as necessary, downsample reads to generate standardized summary data across samples. This process ultimately yields a matrix encompassing gene counts and cells.

All statistical analyses were conducted using R and Python. Cells exhibiting fewer than 500 observed genes were excluded from the analysis. Additionally, cells were eliminated if more than 20% of their reads mapped to mitochondrial genes.

### Unsupervised clustering and identification

The Seurat is used for normalization and cell clustering based on differential gene expression data [56,57]. These corrected data are used for downstream analysis after permutation and selection of the top 50 principal components based on principal component analysis (PCA) scores. Unsupervised clustering was performed in Seurat, where the algorithm uses a graph-based approach to first construct a K-nearest neighbor graph (K=20) and identify clusters by iteratively forming cell groups to optimize the modularization function. The number of clusters was determined using the Louvain algorithm for community detection, which is implemented in Seurat with a resolution of 0.4. The FindMarkers function (test.use=presto) in Seurat was used to select differentially expressed genes (DEGs). The P value <0.05 and |log2foldchange|>0.25 were set as the threshold for significant differential expression.

### Pseudo-time analysis

Use the Monocle R package to determine the developmental pseudo-time [58]. First, use the importCDS function in Monocle to convert the raw counts from the Seurat object to a CellDataSet object. Use the differentialGeneTest function in the Monocle2 package to select the sorted genes (*q* val < 0.01), which may provide information when sorting cells along the pseudo-time trajectory. Use the reduceDimension function for dimensionality reduction clustering analysis, and then use the default parameters for trajectory inference to track changes in pseudo-time.

### CellChat analysis

To perform cell communication analysis using the CellChat R package [59], first we import the standardized expression matrix and create a cellchat object using the createCellChat function. Next, we preprocess the data using default parameters. Then we identify any potential ligand-receptor interactions. Finally, we use the aggregateNet function to summarize the cell communication network.

### Transcription factor analysis

Transcription factor analysis was performed using python-based pySCENIC [60]. The normalized cell transcriptome expression matrix was extracted from R and analyzed using pySCENIC. The results were imported back into R for visualization.

### Gene set enrichment analysis (GSEA)

GSEA was used to complete KEGG term enrichment analysis with the Molecular Signatures Database (MSigDB) C2 KEGG gene sets Version 7.2 separately [61].

### Cell culture

LAD2 cells were cultured in StemPro™-34 SFM medium containing 2mM L-glutamine + 100ng/ml SCF + 1% P/S. WPMY-1 was cultured in DMEM medium containing 10% FBS and 1% P/S.

ECFC cells were cultured from donors’ peripheral blood in EGM-2 medium containing 2mM L-glutamine + 2U/mL heparin sodium+ 10% Human Platelet Lysate + 1% P/S [62]. ECFC showed a typical cobblestone appearance. ECFC within 3 generations were used for experiments.

Primary BPH fibroblasts were derived from the donor’s prostate transition zone tissue. The prostate transition zone tissue was cut into pieces and suspended in HBSS containing 1000 U/mL collagenase I + 1 mg/mL DNase I + 1% P/S, and digested at 37°C for 1 hour. Afterwards, the digested tissue was quickly added to DMEM medium containing 10ng/ml FGF-basic + ITS-G + 10% FBS + 1% P/S for culture. After about 7 days, it was observed that primary fibroblasts crawled out from the adherent tissue pieces. The fibroblasts used in the experiment were all passaged for 3-5 generations.

To avoid mycoplasma contamination, cells were examined by cytoplasmic DAPI staining every three passages. All cells were cultured at 37°C in an environment containing 5% CO2, and cells with a confluence of 60-70% were collected.

### LAD2 cell siRNA transfection

Mix 105μL LAD2 medium (without antibiotics) with 20μL Lipofectamine™ RNAiMAX, and mix 123μL medium with 2μL siRNA (100nM in 2mL). Mix Lipo and siRNA (125μL+125μL), leave at room temperature for 5 minutes, and add 250μL to each well of a six-well plate with LAD2 cells in 1.75mL medium. The following are the siRNA sequences used.

KIT-si-1: CCUUGGAAGUAGUAGAUAAtt

KIT-si-2: CGGUUGAAUGUAAGGCUUAtt

### RT-qPCR

RNA was isolated from cells using Trizol reagent and reverse transcribed using PrimeScript RT reagent according to the manufacturer’s instructions. SYBR green was used for quantitative PCR. The following are the primer sequences used.

*VEGFA*: Forward AGGGCAGAATCATCACGAAGT; Reverse AGGGTCTCGATTGGATGGCA

*TGFB1*: Forward GGCCAGATCCTGTCCAAGC; Reverse GTGGGTTTCCACCATTAGCAC

*KIT*: Forward CGTTCTGCTCCTACTGCTTCG; Reverse CCCACGCGGACTATTAAGTCT

*GAPDH*: Forward GGAGCGAGATCCCTCCAAAAT; Reverse GGCTGTTGTCATACTTCTCATGG

### CUT&Tag library construction

CUT&Tag assay was performed using HyperactiveTM In-Situ ChIP Library Prep Kit for Illumina according to manufacturer’s instruction. Briefly, prepared concanavalin A-coated magnetic beads (ConA beads) were added to resuspended cells and incubated at room temperature to bound cells. Non-ionic detergent Digitonin was used to permeate cell membrane. Then, anti-JUND antibody, secondary antibody and the Hyperactive pA-Tn5 Transposase were incubated with the cells that were bounded by ConA beads in order. Therefore, the Hyperactive pA-Tn5 Transposase can exactly cut off the DNA fragments that were bound with target protein. In addition, the cut DNA fragments can be ligated with P5 and P7 adaptors by Tn5 transposase and the libraries were amplified by PCR with the P5 and P7 primers. The purified PCR products were evaluated using the Agilent 2100 Bioanalyzer. Finally, these libraries were sequenced on the Illumina NovaSeq6000 platform and 150bp paired-end reads were generated for the following analysis.

### CUT&Tag analysis

The raw sequence data were firstly quality trimmed by fatspsoftware to obtain the clean reads. Then the clean reads were aligned to the (RCh38.p12) using Bowtie2 and subsequently analyzed by the SEACR software based the ‘stringent’ parameter to detect genomic regions enriched for multiple overlapping DNA fragments (peaks) that we considered to be putative binding sites. Visualization of peak distribution along genomic regions of interested genes was performed with IGV. Peaks were then annotated using chipseeker software to obtain the genes and gene annotations about peaks.

### Immunohistochemistry

Tissue dewaxing, rehydration, and antigen retrieval steps will be performed. Block endogenous peroxidase activity with 3% hydrogen peroxide. Wash the slides three times with PBS and incubate with primary antibody at room temperature for 2 hours. After washing three times with PBS, apply secondary antibody for 30 minutes at room temperature. Stain with diaminobenzidine, and then counterstain the tissue with hematoxylin.

### Immunofluorescence

Tissue dewaxing, rehydration, and antigen retrieval steps will be performed. 3% hydrogen peroxide is used to block endogenous peroxidase activity. Wash the slides three times with PBS and incubate with primary antibody at room temperature for 2 hours. After washing three times with PBS, incubate the slides with fluorescent secondary antibody at room temperature for 1 hour. Then wash the samples with PBS and treat with DAPI for 3 minutes. Use a high-sensitivity confocal laser microscope (Leica DMI-8) to obtain micrographs.

### RNAscope

According to the manufacturer’s instructions, RNAscope was performed using the following probes and reagents. In short, before the fluorescently labeled RNAscope hiplex probes hybridize with the target RNA, the paraffin tissue sections undergo dewaxing treatment and signal amplification using the RNAscope Hiplex v2 reagent during multiple rounds of probe hybridization and stripping. Representative images were taken using a Leica DMI-8 confocal system, and the ROI position was stored for repositioning during multiple rounds of hybridization. After stripping and imaging, the Leica LAS X software was used to remove the surrounding fluorescence by using a reference image of the same ROI without probes.

### Multiplexed immunofluorescence staining

Samples were cut into 5 μm thick sections and loaded onto adhesion microscope slides. The slides were preprocessed with deparaffinization, rehydration, and antigen retrieval for mIHC staining. Multiplexed immunofluorescence staining of tissue was performed using TG TSA Multiplex IHC Assay Kits.

### Western blot

5-20 μL of cell lysate was boiled in 1× loading buffer for 7 minutes and loaded onto a sodium dodecyl sulfate-polyacrylamide gel electrophoresis (SDS-PAGE) gel. After electrophoresis, the proteins were transferred to a polyvinylidene fluoride membrane, which was blocked with Tris-buffered saline (TBST) containing 5% skim milk and Tween 20 at room temperature for 2 hours. The membrane was then incubated overnight with diluted primary antibody in 5% bovine serum albumin (BSA) at 4°C, washed with TBST at room temperature, and incubated with corresponding secondary antibody for 2 hours at room temperature.

### In *vivo* assay

All animal experiments were approved by the Animal Ethics Committee of the Animal Experimental Center of Shanghai General Hospital. All mice were raised under specific pathogen-free (SPF) conditions. All experiments used 8-week-old mice. A BPH model was established by subcutaneously injecting 5 mg/kg testosterone propionate (dissolved in sterile corn oil) for 28 consecutive days.

In the mast cell experimental group, the control group mice were injected with sterile corn oil subcutaneously daily, while the remaining mice were injected with 5 mg/kg/d testosterone propionate subcutaneously for 28 consecutive days. In the drug treatment experimental group, the control group mice were injected with sterile corn oil subcutaneously daily, while also given 0.4 ml of 0.5% CMC-Na by gavage. The BPH modeling group mice were injected with 5 mg/kg testosterone propionate subcutaneously daily, while also given 0.4 ml of 0.5% CMC-Na by gavage. The mice treated with drugs were all injected with 5 mg/kg testosterone propionate subcutaneously daily, and were given 50 mg/kg/d imatinib, 30 mg/kg/d sunitinib, 30 mg/kg/d cabozantinib, 30 mg/kg/d regorafenib, 100 mg/kg/d tranilast, 20 mg/kg/d sodium cromoglicate, 15 mg/kg/d ketotifen, and 30 mg/kg/d asiaticoside, all dissolved in 0.4 ml of 0.5% CMC-Na and given to the mice by gavage. The drug treatment experiment lasted for 28 days.

After the planned administration period ended, the mice were euthanized with CO2, weighed, and the bladder and part of the urethra were removed. The mouse prostate was separated using a stereomicroscope, weighed, and fixed with 4% paraformaldehyde. The mouse prostate index = mouse prostate weight (mg) / mouse body weight (g).

### statistical analysis

Data are expressed as mean ± SEM. All tissue section image statistics were completed using Fiji [63]. Differences between groups were analyzed using Student’s t test or Chi-square test, depending on the sample type. Statistical analysis was performed using Graphpad Prism 10.2. The *p* value less than 0.05 was considered significant. In the data graphs, significance was indicated by ns *p*≥0.05, \**p*<0.05, \*\**p*<0.01, \*\*\**p*<0.001, and \*\*\*\**p*<0.0001.

### Data availability

The raw sequence data of samples L1-L3 and S1-S3 reported in this paper have been deposited in GEO under the accession number GSE226237; the raw sequence data of samples L4-L6 and S4-S5 reported in this paper have been deposited in the Genome Sequence Archive (Genomics, Proteomics & Bioinformatics 2021) in National Genomics Data Center (Nucleic Acids Res 2024), China National Center for Bioinformation / Beijing Institute of Genomics, Chinese Academy of Sciences (GSA-Human: HRA009899) that are publicly accessible at https://ngdc.cncb.ac.cn/gsa-human.

### Other key resources

Other key resources are listed in Table S1 (key resources table).

## Acknowledgments

We would like to express our gratitude to Jia Li, Juan Wang, Lihui Lin, Xia Peng from the Department of Laboratory Medicine of Shanghai General Hospital for providing the equipment required for our experiments. We thank Yuji Huang from the Department of Laboratory Medicine of Shanghai General Hospital for his guidance on our LAD2 cell siRNA transfection experiment. We thank Tiewen Li and Zheng Zhou from the Department of Urology of Shanghai General Hospital for their assistance with our animal experiments. We would also like to thank Feng Xie from the Shanghai Institute of Immunology, Shanghai Jiao Tong University School of Medicine for her advise of this research. This research was funded by a grant from National Natural Science Foundation of China (No. 82270810 and No. 81871267).

## Author Contributions

TC, DZ, LQ, BM, CY, LL and YJ conceived the idea of the study; JT, TC, DZ, CF and YS performed the bioinformatics analyses; TC, JX, YL, MW, WC and HL performed the animal experiment; WG and JH were involved in acquisition of patient tissues and performed pathological analyses; TC, JX and YL conducted cytological experiments; DC, YZ, XW and JL were involved in acquisition of patient tissues or accessing medical records; YR, BH and SX contributed in the technical, material, or administrative support of the study. all authors discussed the results and revised the manuscript.

## Conflict of Interest

The authors declare that there are no conflicts of interest.

## Notes

### Competing Interest Statement

The authors have declared no competing interest.

