## Supplementary material for "Senescent mast cells contribute to the progression of benign prostatic hyperplasia via SCF/c-KIT mediated endothelial-mesenchymal transition": Figure S1-S15, Table S1

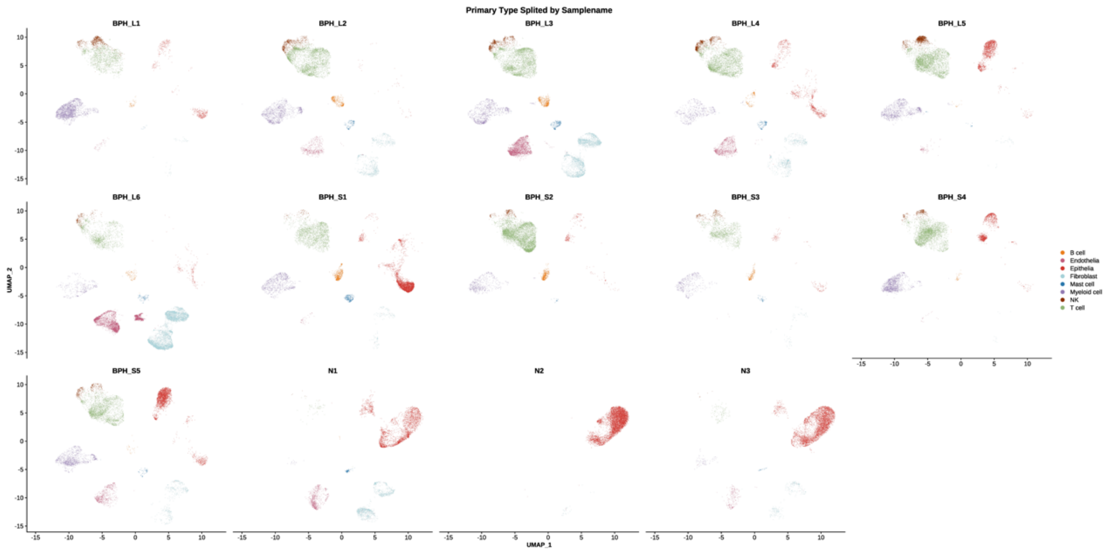


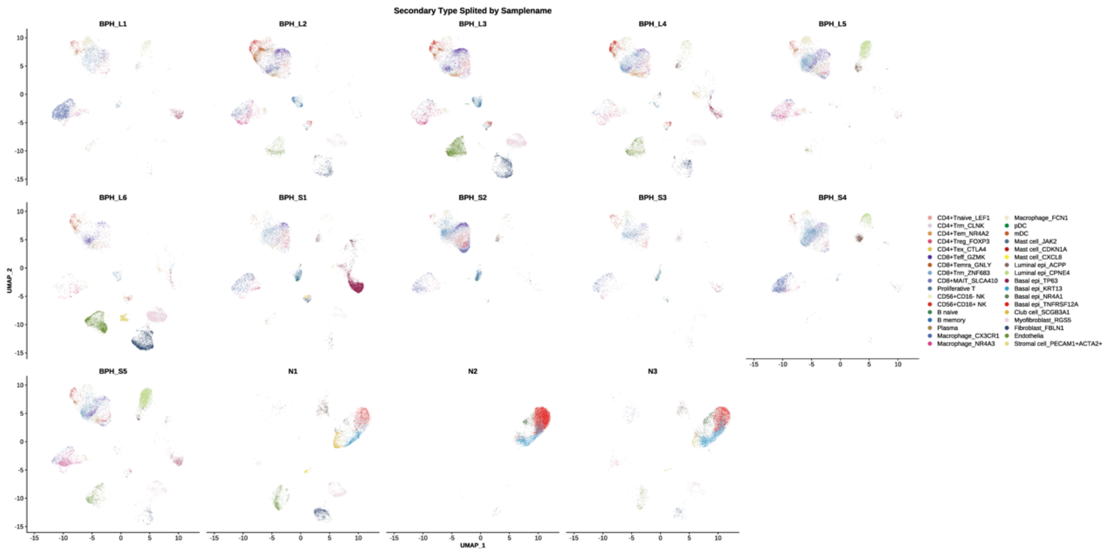


**Figure S1-S2**

UMAP plot of prostate cell subsets with sample-specific color annotation.

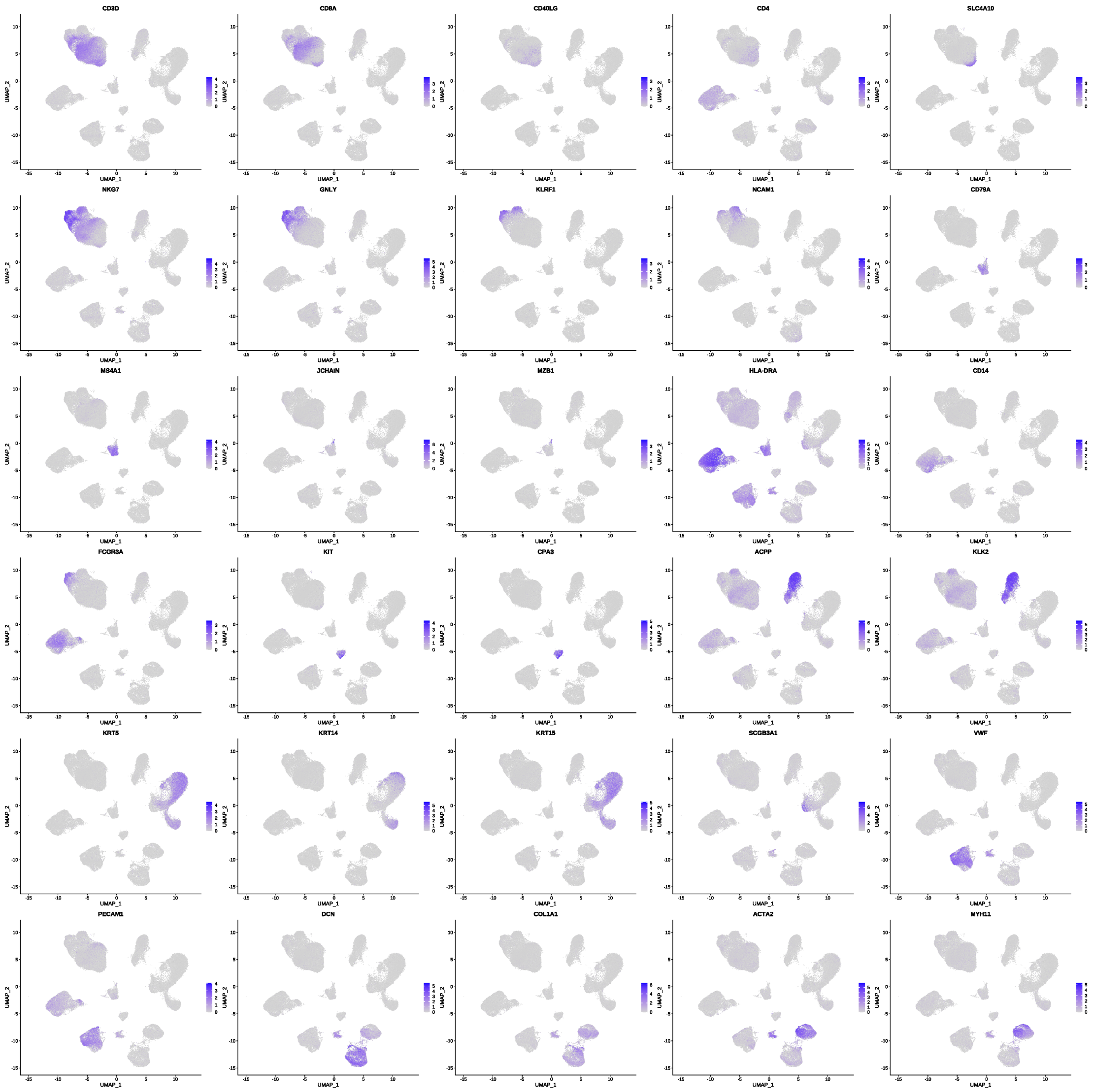


**Figure S3**

UMAP plots showing the distribution of expression of selected signature genes for all cell subsets.

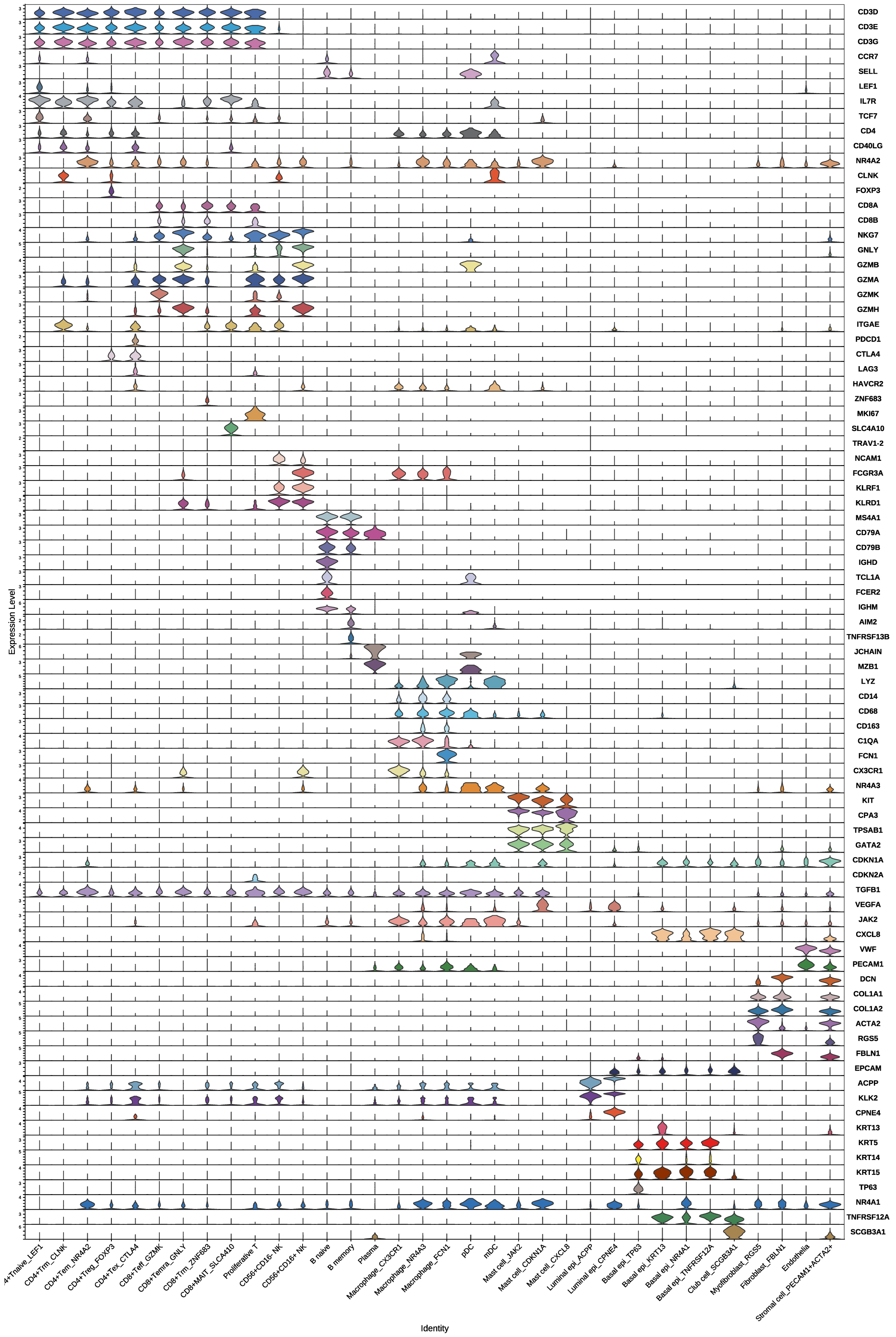
Expression Level
CD3D
CD3E
CD3G
CCR7
SELL
LEF1
IL7R
TCF7
CD4
CD40LG
NR4A2
CLNK
FOXP3
CD8A
CD8B
NKG7
GNLY
GZMB
GZMA
GZMK
GZMH
ITGAE
PDCD1
CTLA4
LAG3
HAVCR2
ZNF683
MKI67
SLC4A10
TRAV1-2
NCAM1
FCGR3A
KLRF1
KLRD1
MS4A1
CD79A
CD79B
IGHD
TCL1A
FCER2
IGHM
AIM2
TNFRSF13B
JCHAIN
MZB1
LYZ
CD14
CD68
CD163
C1QA
FCN1
CX3CR1
NR4A3
KIT
CPA3
TPSAB1
GATA2
CDKN1A
CDKN2A
TGFB1
VEGFA
JAK2
CXCL8
VWF
PECAM1
DCN
COL1A1
COL1A2
ACTA2
RGS5
FBLN1
EPCAM
ACPP
KLK2
CPNE4
KRT13
KRT5
KRT14
KRT15
TP63
NR4A1
TNFRSF12A
SCGB3A1
4+Tnaive_LEF1
CD4+Trm_CLNK
CD4+Tem_NR4A2
CD4+Treg_FOXP3
CD4+Tex_CTLA4
CD8+Teff_GZMK
CD8+Temra_GNLY
CD8+Trm_ZNF683
CD8+MAIT_SLCA410
Proliferative T
CD56+CD16- NK
CD56+CD16+ NK
B naive
B memory
Plasma
Macrophage_CX3CR1
Macrophage_NR4A3
Macrophage_FCN1
pDC
mDC
Mast cell_JAK2
Mast cell_CDKN1A
Mast cell_CXCL8
Luminal epi_ACPP
Luminal epi_CPNE4
Basal epi_TP63
Basal epi_KRT13
Basal epi_NR4A1
Basal epi_TNFRSF12A
Club cell_SCGB3A1
Myofibroblast_RGS5
Fibroblast_FBLN1
Endothelia
Stromal cell_PECAM1+ACTA2+
Identity

**Figure S4**

Violin plots showing the distribution of expression of selected signature genes for 34 cell subsets.

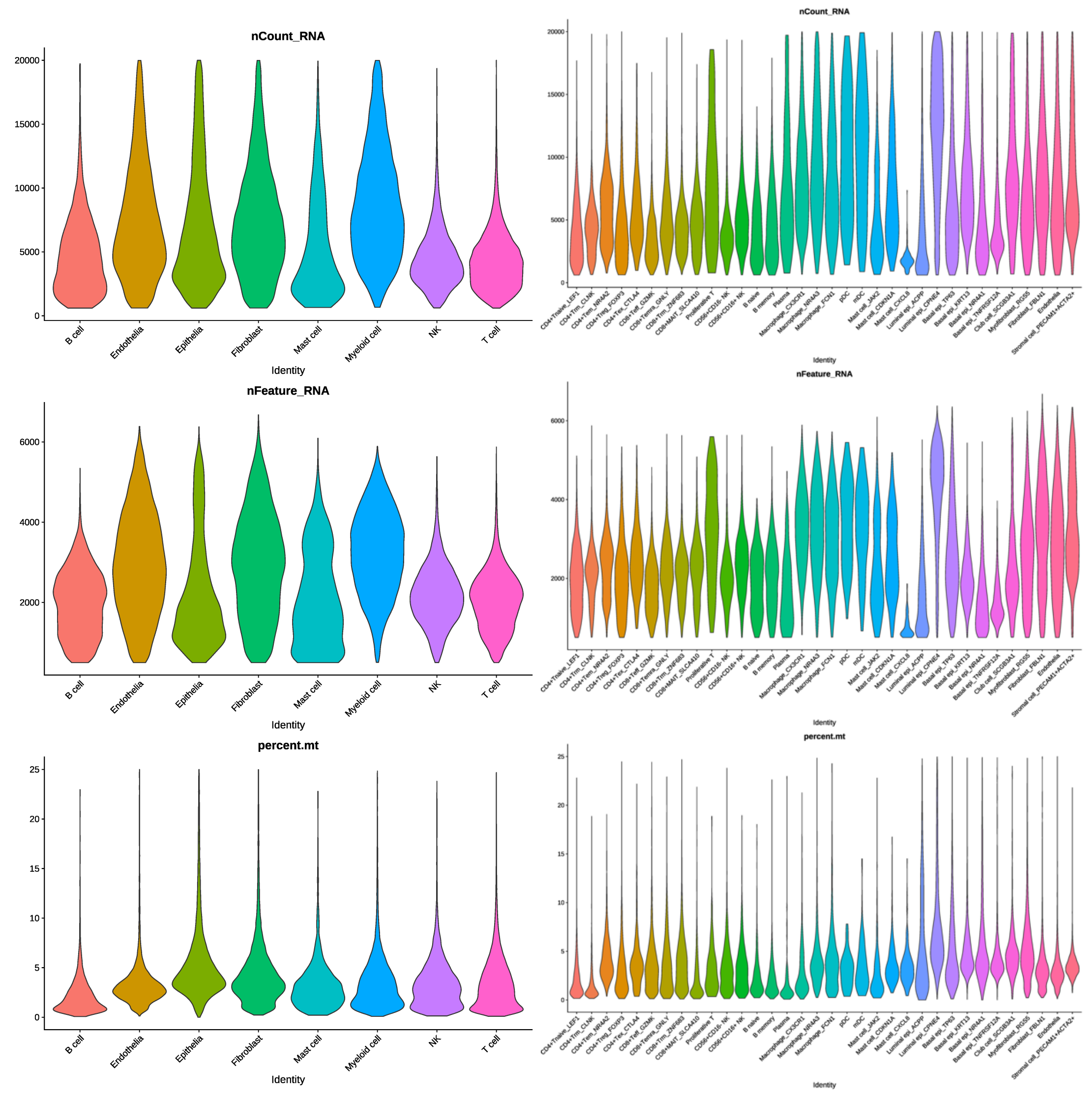


**Figure S5-S6**

Quality control metric distributions across prostate cell types

Violin plots display UMI counts (transcript abundance), detected genes (complexity), and mitochondrial gene percentage (cell viability) per annotated lineage.

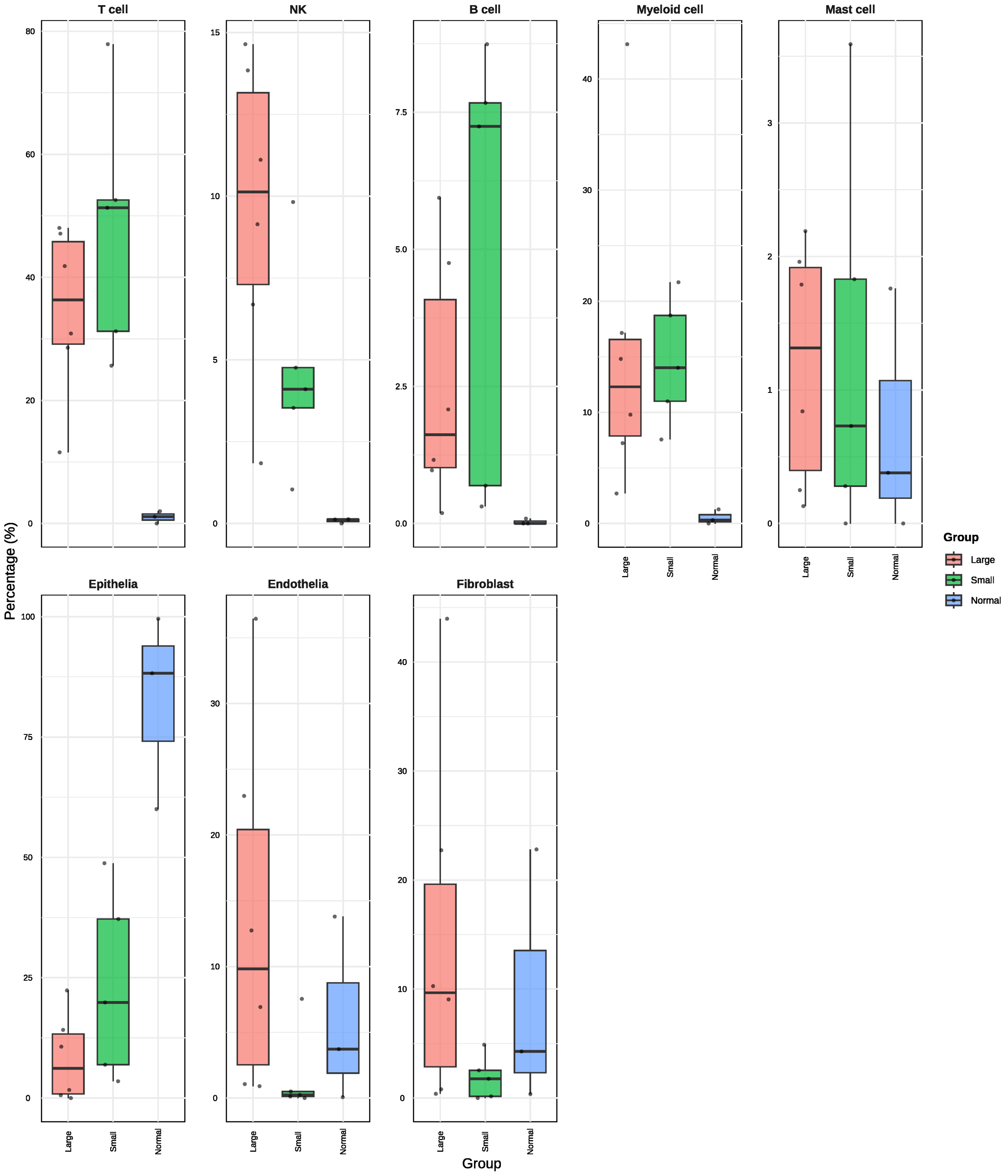


**Figure S7**

Stacked bar plot quantifying the proportional distribution of annotated cell types across experimental groups

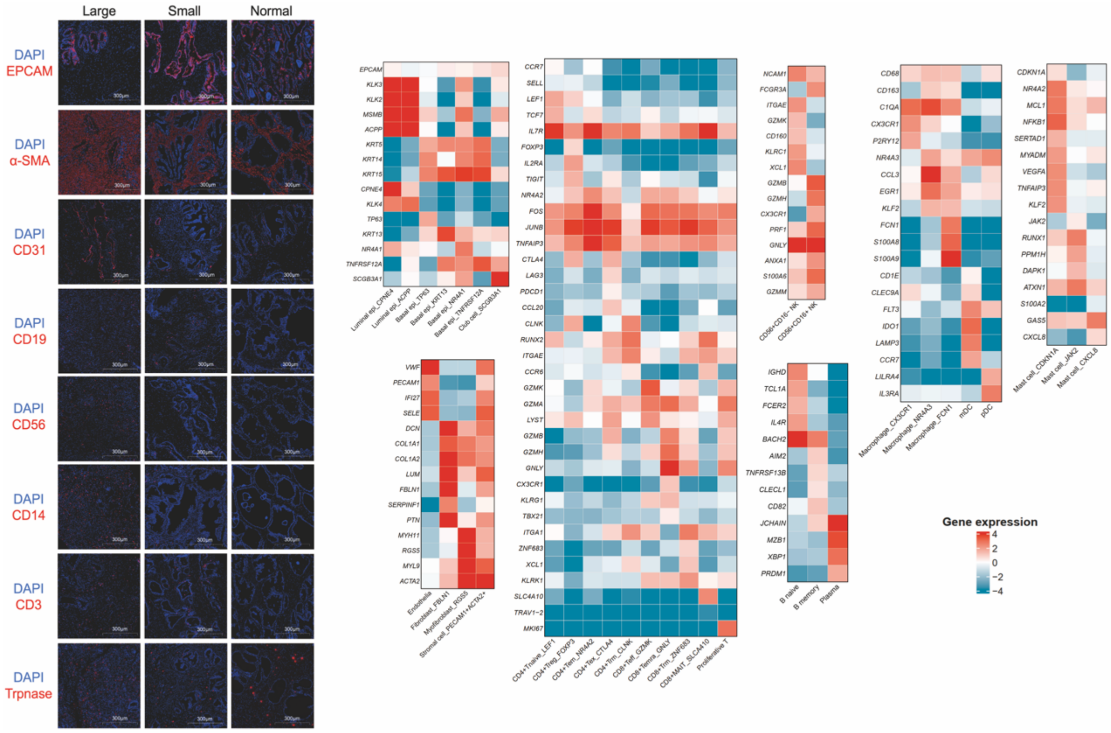


**Figure S8**

(A) Immunofluorescence staining shows celltype in different experimental groups of human prostate. Epithelial cells (EPCAM), endothelial cells (CD31), stromal cells (α-SMA), T cells (CD3), B cells (CD19), NK cells (CD56) and mast cells (Trpnase).

(B) Heatmap displaying lineage-defining gene expression signatures across each cell subsets.

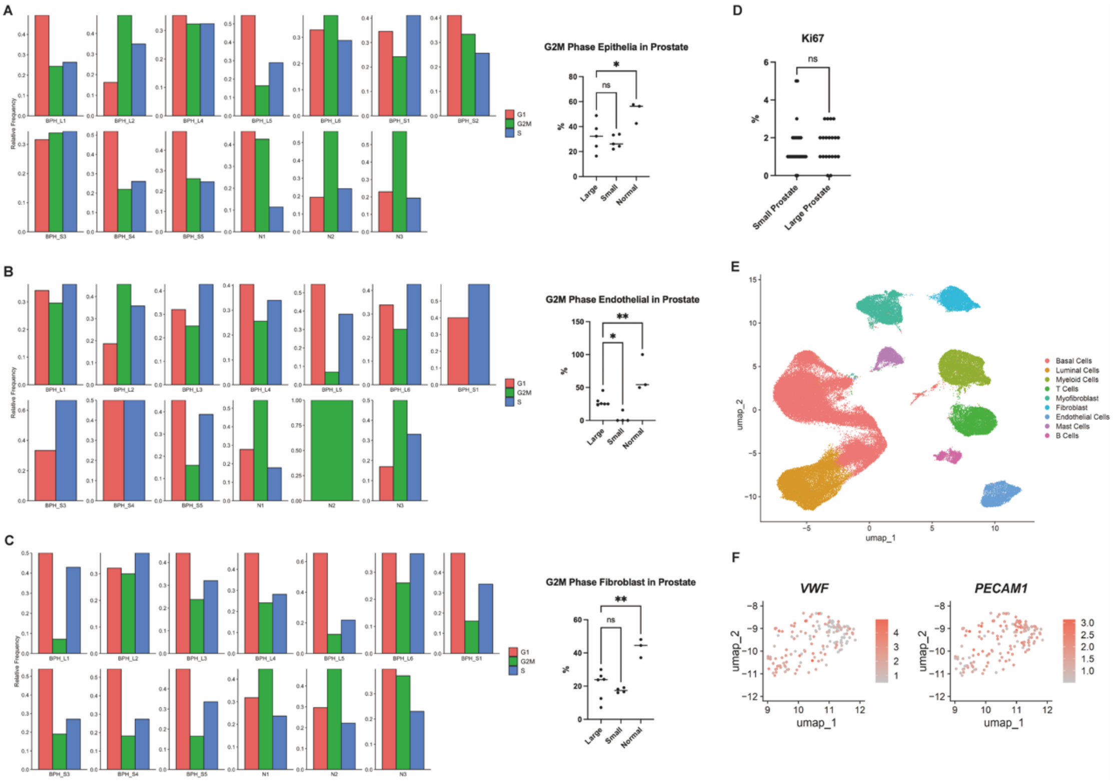


**Figure S9**

(A) The proportion of epithelia cell cell cycle and the proportion of G2M phase cells in each sample.

(B) The proportion of endothelial cell cycle and the proportion of G2M phase cells in each sample.

(C) The proportion of fibroblast cycle and the proportion of G2M phase cells in each sample.

(D) The statistical situation of Ki67 expression in large and small prostates. (Small, n=26; large, n=20)

(E) UMAP plot of annotated cell types in the human prostate (dataset from GSE145928 and GSE183676).

(F) Feature plot visualizing expression of *VWF* and *PECAM* in the human prostate (dataset from GSE145928 and GSE183676).

50

### Figure S10

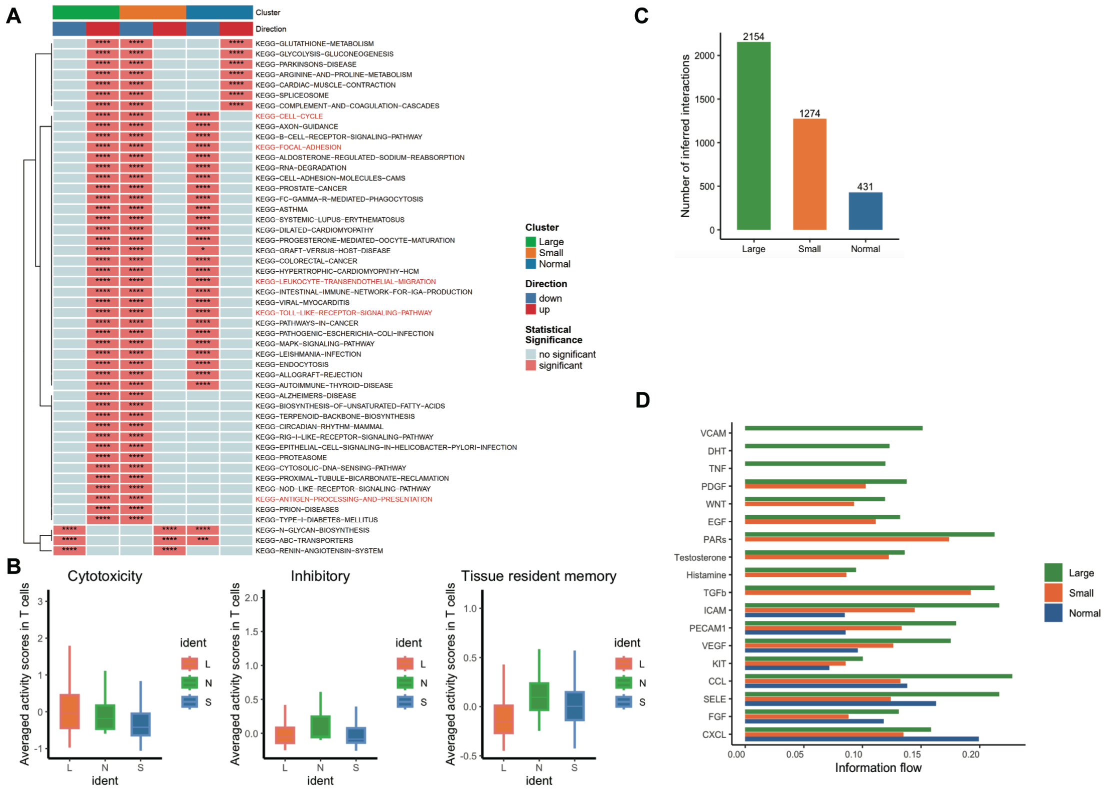


(A) Functional enrichment heatmap of T cell DEGs across experimental groups, analyzed by irGSEA (ssGSEA method, FDR < 0.05).

(B) The bar chart displaying T cell activity scores.

(C) Bar plot summarizing interaction frequencies between major cell clusters across experimental groups.

(D) Bar plot of each pathway between major cell clusters across experimental groups.

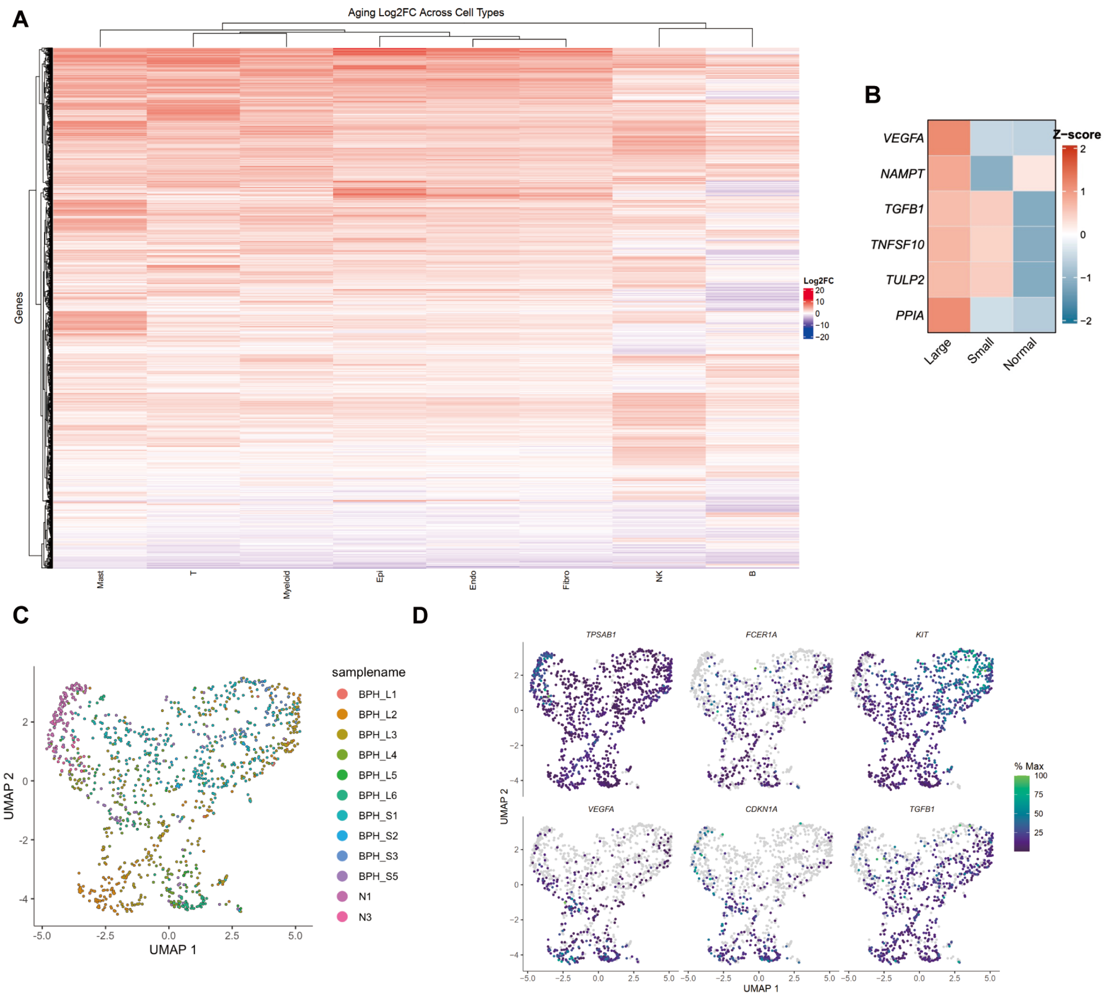


**Figure S11**

(A) Heatmap displaying log2FCs in gene expression (aged versus young) of human prostate aging-associated DEGs in each cell type.

(B) The heatmap reveals the differential expression of mast cell secretory factors in the prostates of young individuals classified as large, small, and normal-sized.

(C) UMAP plot of mast cell using Monocle3-based dimensionality reduction and clustering

(D) Feature plot visualizing specific genes expression of mast cells in the prostate.

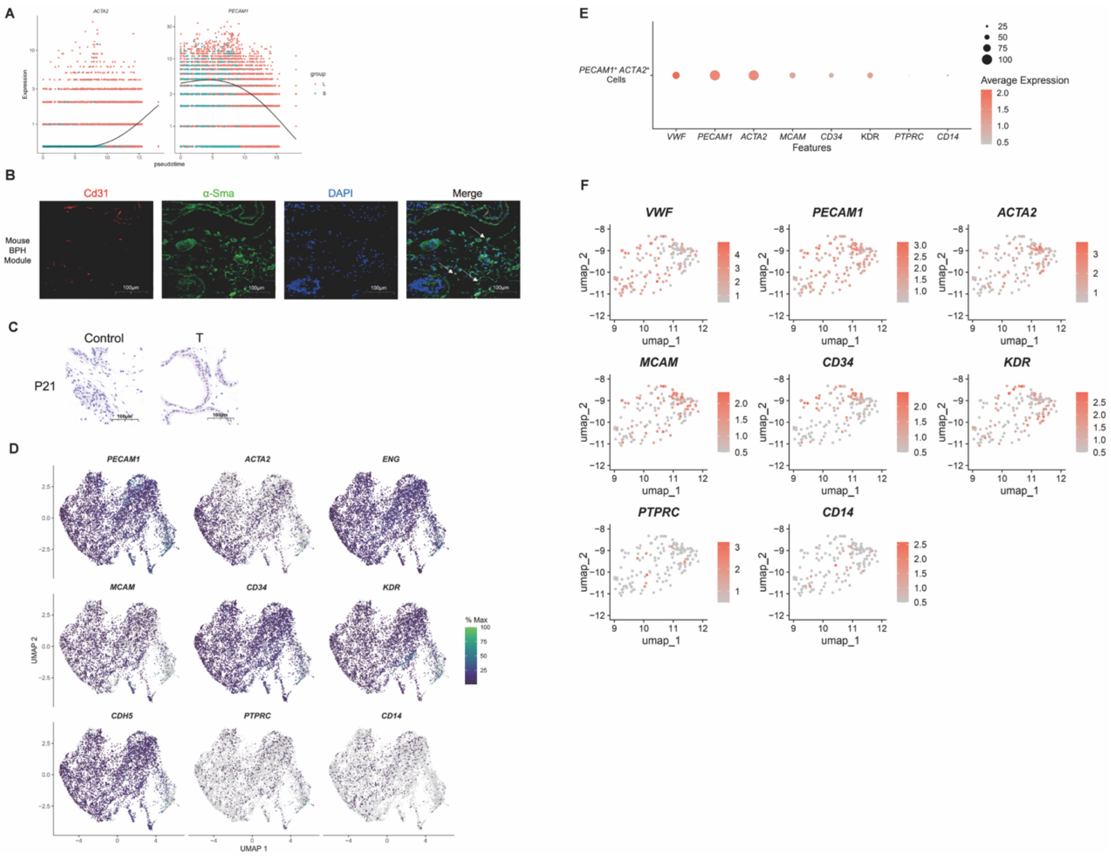


**Figure S12**

(A) Pseudo-time analysis reveals the changes in *PECAM1* and *ACTA2*.

(B) Multicolor immunohistochemistry reveals the presence of Cd31+ α-Sma+ cells in mouse BPH tissue induced by testosterone propionate.

(C) Immunohistochemistry staining shows the expression of P21 in different experimental groups of mouse prostate.

(D) Feature plot characterizing the expression of ECFC-specific genes in endothelial cells.

(E) Dot plot demonstrating expression of key marker genes in *PECAM1+ACTA2+* stromal cell (dataset from GSE145928 and GSE183676).

(F) Feature plot demonstrating expression of key marker genes in *PECAM1+ACTA2+* stromal cell (dataset from GSE145928 and GSE183676).

85

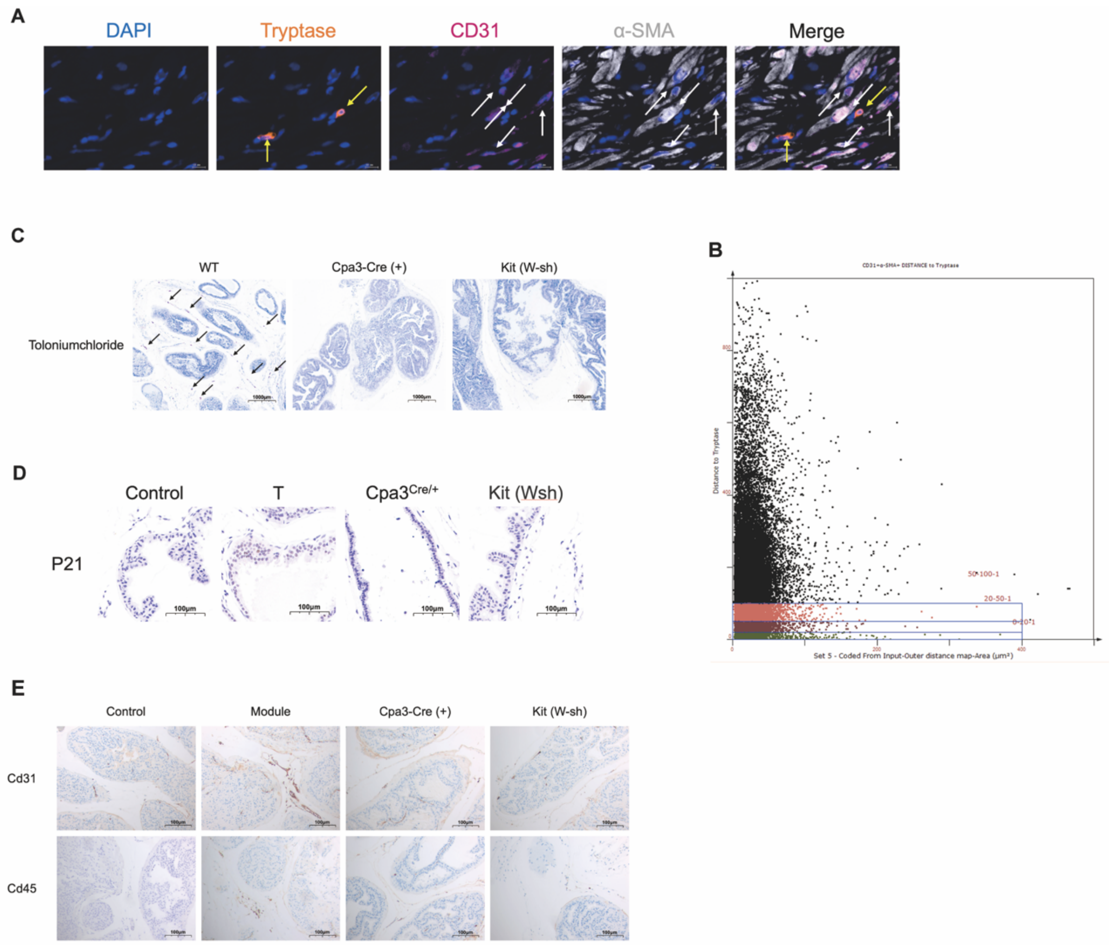


**Figure S13**

(A-B) TissueFAXS Spectra reveals the spatial relationship between mast cells (Tryptase+ cells) and CD31+ α-SMA + cells.

(C) Toloniumchloride staining confirmed that there were no mast cells in mast cell knockout mice.

(D-E) Immunohistochemical staining was performed on mouse prostate tissue using anti-P21, anti-Cd31 and anti-Cd45 antibodies to show the vascular density and immune cell infiltration in the prostate tissue.

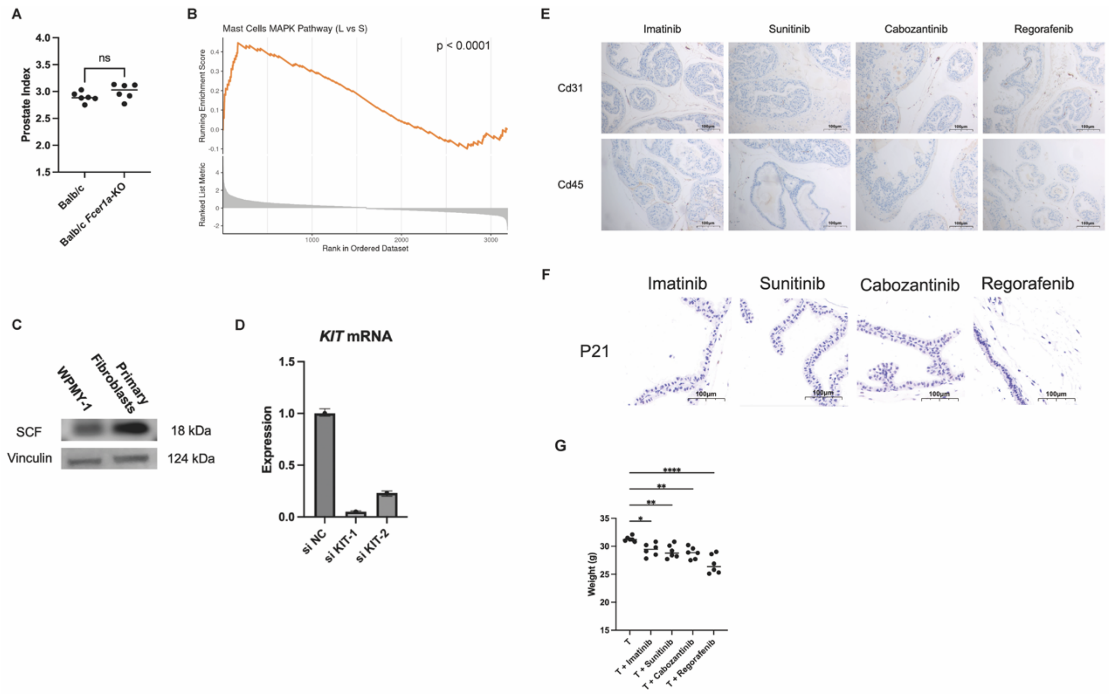


**Figure S14**

(A) Prostate index of mice in the model group, and *Fcer1a*-KO group. All mice received testosterone propionate subcutaneously at 5 mg/kg/d.

(B) GSEA analysis of transcriptome sequencing data showed that the MAPK pathway was up-regulated in mast cells in large prostates.

(C) Western-blot revealed that primary fibroblasts derived from the large prostate expressed SCF.

(D) qPCR revealed the efficiency of si-KIT knockdown in mast cells.

(E-F) Immunohistochemical staining was performed on mouse prostate tissue using anti-P21, anti-Cd31 and anti-Cd45 antibodies to show the vascular density and immune cell infiltration in the prostate tissue.

(G) Weight of mice in the control group, model group, and TKIs-treated group.

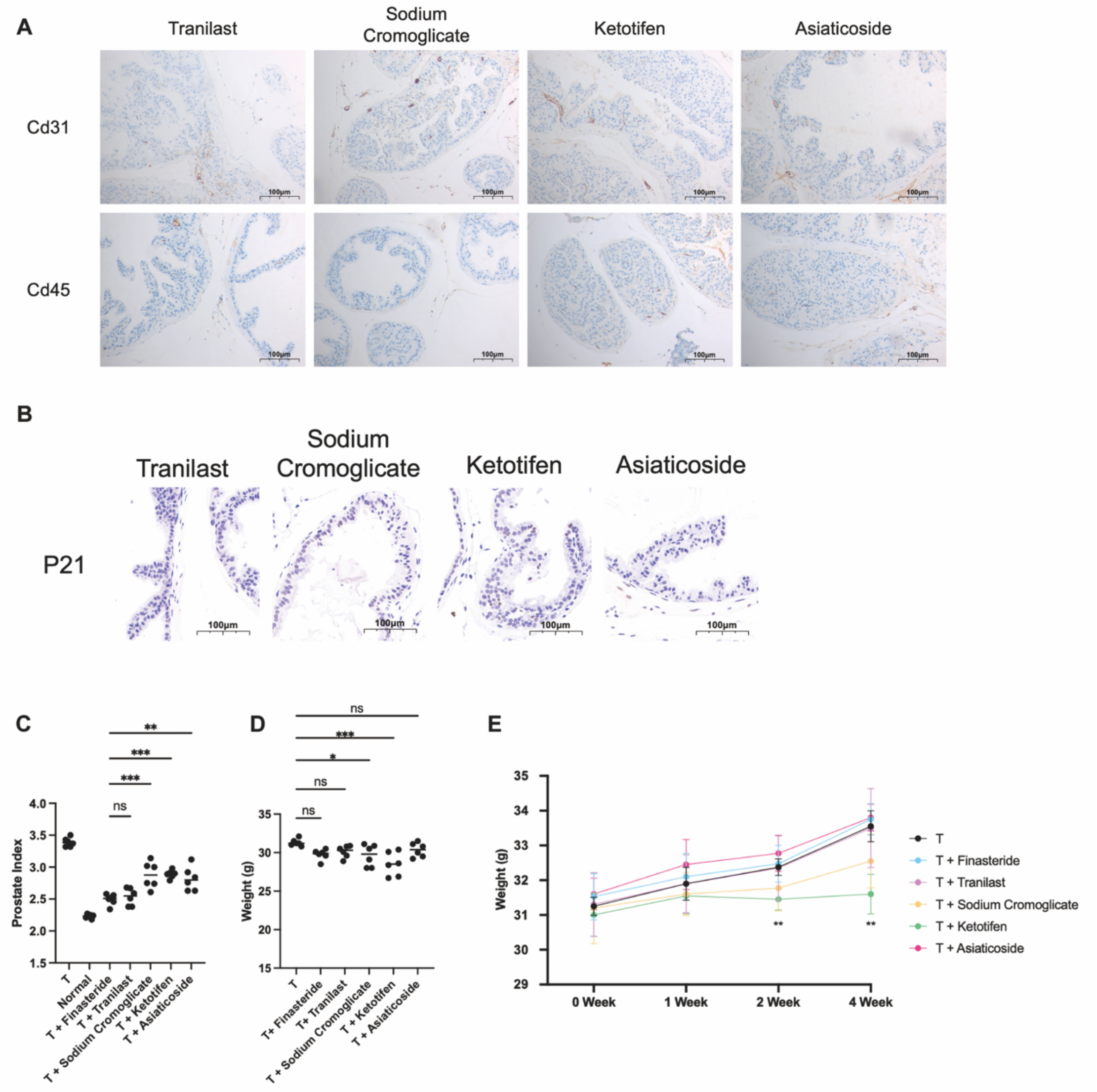


**Figure S15**

(A-B) Immunohistochemical staining was performed on mouse prostate tissue using anti-P21, anti-Cd31 and anti-Cd45 antibodies to show the vascular density and immune cell infiltration in the prostate tissue.

(C) Prostate index of mice in different treatment group. T stands for daily subcutaneous injection of 5mg/kg/d testosterone propionate.

(D-E) Weight of mice in the control group, model group, and mice prevented/treated with mast cell membrane stabilizers and TGF-β inhibitors.

| Table S1 Key resources table | | |
|---|---|---|
| **REAGENT or RESOURCE** | **SOURCE** | **IDENTIFIER** |
| **Antibodies** | | |
| Anti-alpha smooth muscle Actin | Abcam | Cat# ab7817; RRID:AB_262054 |
| Anti-EPCAM | Abcam | Cat# ab223582; RRID:AB_2762366 |
| Anti-CD3 | Abcam | Cat# ab11089; RRID:AB_2889189 |
| Anti-CD19 | Abcam | Cat# ab134114; RRID:AB_2801636 |
| Anti-CD56 (NCAM1) | Abcam | Cat# ab220360; RRID:AB_2927664 |
| Anti-CD14 | Abcam | Cat# ab2378; RRID:AB_303023 |
| Anti-Trpnase | Abcam | Cat# ab183322; RRID:AB_2909464 |
| Anti-p21 (human) | Abcam | Cat# ab109520; RRID:AB_1086053 7 |

| | | |
|---|---|---|
| Anti-p21 (mouse) | Abcam | Cat# ab188224; RRID:AB_2734729 |
| Anti-CD31 | Proteintech | Cat# 11265-1-AP; RRID:AB_2299349 |
| Anti-c-Kit | Abcam | Cat# ab32363; RRID:AB_731513 |
| Anti-Beta Tubulin (HRP conjugated) | Proteintech | Cat# HRP-66240; RRID:AB_2883838 |
| Anti-ERK1/2 | Proteintech | Cat# 11257-1-AP; RRID:AB_2139822 |
| Anti-Phospho-p44/42 MAPK (Erk1/2) | CST | Cat# 4370; AB_2315112 |
| Anti-Phospho-c-Kit | CST | Cat# 3073; RRID:AB_1147635 |
| Anti-Vinculin (HRP conjugated) | CST | Cat# 18799; RRID:AB_2714181 |
| Anti-CD45 | Invitrogen | Cat# 14-0451-82; RRID:AB_467251 |
| Anti-SCF | Proteintech | Cat# 26582-1-AP; RRID:AB_2880561 |

| | | |
|---|---|---|
| Anti-CD34 | Abcam | Cat# ab81289; RRID:AB_1640331 |
| Anti-CD146 | Abcam | Cat# ab75769; RRID:AB_2143375 |
| Anti-VEGFR2 | Abcam | Cat# ab115805; RRID:AB_10899278 |
| Anti-JUND | Active Motif | Cat# 61403; RRID:AB_2793621 |
| **Chemicals, and recombinant proteins** | | |
| DMEM, no phenol red | Gibco | Cat# 21063029 |
| Fetal Bovine Serum | Gibco | Cat# 10099141C |
| StemPro™-34 SFM (1X) | Gibco | Cat# 10639011 |
| EGM-2 Endothelial Cell Culture Medium | Lonza | Cat# CC-3162 |
| Human Platelet Lysate | CytoFamily | Cat# C8010-0100 |
| Heparin Sodium | Absin | Cat# abs47014863 |

| | | |
|---|---|---|
| ITS -G | Gibco | Cat# 41400045 |
| L-glutamine | Gibco | Cat# 25030081 |
| Collagenase I | Biosharp | Cat# BS163 |
| DNase I | Absin | Cat# abs47047435 |
| TrypLE Express | Gibco | Cat# 12605028 |
| Recombinant Human FGF-basic | PeproTech | Cat# 100-18B |
| Recombinant Human SCF | PeproTech | Cat# 300-07 |
| Lipofectamine™ RNAiMAX | Invitrogen | Cat# 13778150 |
| PrimeScript™ RT Master Mix | Takara | Cat# RR036 |
| qPCR SYBR Green | Yeasen | Cat#11203ES |
| Hanks' Balanced Salt Solution (with Ca2+ & Mg2+) | Beyotime | Cat# C0219 |
| Testosterone Propionate | MCE | Cat# HY-B1269 |
| Corn oil (Sterile) | Beyotime | Cat# ST2308 |
| DMSO | Beyotime | Cat# ST038 |

| | | |
|---|---|---|
| PD98059 | MCE | Cat# HY-12028 |
| Imatinib | MCE | Cat# HY-15463 |
| Sunitinib | MCE | Cat# HY-10255A |
| Cabozantinib | MCE | Cat# HY-13016 |
| Regorafenib | MCE | Cat# HY-10331 |
| Tranilast | MCE | Cat# HY-B0195 |
| Cromolyn sodium | MCE | Cat# HY-B0320A |
| Ketotifen fumarate | MCE | Cat# HY-B0157A |
| Asiaticoside | MCE | Cat# HY-N0439 |
| **Critical commercial assays** | | |
| TGF-β1 ELISA Kit | Multi Science | Cat# EK981 |
| VEGFA ELISA Kit | Abcam | Cat# ab119566 |
| RNAscope HiPlex12 Reagents Kit (488, 550, 650) v2 | ACD | Cat# 324419 |
| RNAscope™ HiPlex Probe- Hs-ACTA2-O4-T4 | ACD | Cat# 1083431-T4 |
| RNAscope™ Hiplex Probe- Hs-VEGFR2-T5 | ACD | Cat# 312121-T5 |

| | | |
|---|---|---|
| RNAscope™ HiPlex Probe- Hs-MCAM-T7 | ACD | Cat# 601731-T7 |
| RNAscope™ Hiplex Probe- Hs-Cpa3-T8 | ACD | Cat# 486731-T8 |
| RNAscope™ HiPlex Probe- Hs-PTPRC-T9 | ACD | Cat# 601991-T9 |
| RNAscope™ HiPlex Probe- Hs-CD34-T10 | ACD | Cat# 560821-T10 |
| RNAscope™ HiPlex Probe- Hs-CD14-T11 | ACD | Cat# 418801-T11 |
| RNAscope™ HiPlex Probe- Hs-PECAM1-O1-T12 | ACD | Cat# 487381-T12 |
| RNAscope™ HiPlex12 Negative Control Probe | ACD | Cat# 324341 |

| | | |
|---|---|---|
| RNAscope™ HiPlex12 Positive Control Probe – Hs | ACD | Cat# 324311 |
| RNAscope™ HiPlex Probe Diluent | ACD | Cat# 324301 |
| TG TSA Multiplex IHC Assay Kits | TG | N/A |
| **Experimental Models: Cell Lines** | | |
| LAD2 | This paper | N/A |
| ECFC | This paper | N/A |
| Prostate Fibroblast | This paper | N/A |
| **Experimental Models: Organisms/Strains** | | |
| C57BL/6J Mouse | Jackson Laboratory | Stock No: 000664 |
| BALB/c Mouse | Jackson Laboratory | Stock No: 000651 |
| ROSA26-LSL-tdTomato Mouse | Cyagen | Stock No: C001476 |
| Cdh5-CreERT2 Mouse | Cyagen | Stock No: C001481 |
| Cpa3-Cre Mouse | Viewsolid | Stock No: VSM30020 |

| | | |
|---|---|---|
| Kit (w-sh) Mouse | Jackson Laboratory | Stock No: 030764 |
| Fcer1a-KO mouse | Viewsolid | N/A |
| **Software and Algorithms** | | |
| Prism 10.2 | https://www.graphpad.com | Commercial |
| Leica LAS X 1.4.6 | https://www.leica-microsystems.com | Commercial |
| Python | https://www.python.org/ | Commercial |
| Fiji | https://imagej.net/software/fiji | Schindelin et al. [63] |
| Cell Ranger 7.0 | https://www.10xgenomics.com/support/software/cell-ranger | Commercial |
| R 4.2.3 | https://cran.r-project.org | Commercial |
| RStudio Server 2023.03.0 Build 386 | https://posit.co/products/open-source/rstudio | Commercial |
| Seurat 5.0.3 | https://satijalab.org/seurat | Hao Y, et al. [56] |
| Seurat 4.3.0 | https://satijalab.org/seurat | Hao Y, et al. [57] |
| monocle3 1.3.3 | https://cole-trapnell-lab.github.io/monocle3/ | Trapnell C, et al. [58] |
| Cellchat 2.1.2 | https://github.com/sqjin/CellChat | Jin S, et al. [59] |
| pySCENIC 0.12.1 | https://github.com/aertslab/pySCENIC | Aibar S, et al. [60] |
| GSEA 3.0 | https://www.gsea-msigdb.org/gsea/index.jsp | Subramanian A, et al. [61] |
| BioRender | https://www.biorender.com | Commercial |

### LAD2 Cell Line Authentication

NIH Provider MTA Modified

**Simple Letter Agreement for the Transfer of Materials**

*[see Federal Register. Dec. 23, 1999, Vol. 64, No 246 [Notices] pp. 72090-72096.]*

**2012-0904**

Page 1 of 2

In response to the RECIPIENT's request for the MATERIAL[*insert description*]

LAD2 cells

*[Are the MATERIALS of human origin?* ☒ *Yes* ☐ *No. If Yes, were Research Materials collected according to 45 CFR Part 46, "Protection of Human Subjects"?* ☒ *Yes* ☐ *No. If Yes, please provide Assurance Number:* *00005897*.]

the PROVIDER asks that the RECIPIENT and the RECIPIENT SCIENTIST agree to the following before the RECIPIENT receives the MATERIAL:

1. The above MATERIAL is the property of the PROVIDER and is made available as a service to the research community.
2. THIS MATERIAL IS NOT FOR USE IN HUMAN SUBJECTS.
3. The MATERIAL will be used for teaching or not-for-profit research purposes only.
4. The MATERIAL will not be further distributed to others without the PROVIDER's written consent. The RECIPIENT shall refer any request for the MATERIAL to the PROVIDER. To the extent supplies are available, the PROVIDER or the PROVIDER SCIENTIST agree to make the MATERIAL available, under a separate Simple Letter Agreement to other scientists for teaching or not-for-profit research purposes only.
5. The RECIPIENT agrees to acknowledge the source of the MATERIAL in any publications reporting use of it.
6. Any MATERIAL delivered pursuant to this Agreement is understood to be experimental in nature and may have hazardous properties. THE PROVIDER MAKES NO REPRESENTATIONS AND EXTENDS NO WARRANTIES OF ANY KIND, EITHER EXPRESSED OR IMPLIED. THERE ARE NO EXPRESS OR IMPLIED WARRANTIES OF MERCHANTABILITY OR FITNESS FOR A PARTICULAR PURPOSE, OR THAT THE USE OF THE MATERIAL WILL NOT INFRINGE ANY PATENT, COPYRIGHT, TRADEMARK, OR OTHER PROPRIETARY RIGHTS. Unless prohibited by law, the RECIPIENT assumes all liability for claims for damages against it by third parties which may arise from the use, storage or disposal of the MATERIAL except that, to the extent permitted by law, the PROVIDER shall be liable to the RECIPIENT when the damage is caused by the gross negligence or willful misconduct of the PROVIDER.
7. The RECIPIENT agrees to use the MATERIAL in compliance with all applicable statutes and regulations.
8. The MATERIAL is provided at no cost, or with an optional transmittal fee solely to reimburse the PROVIDER for its preparation and distribution costs. If a fee is requested, the amount will be indicated here: [*insert fee*] **none**.

The PROVIDER, RECIPIENT and RECIPIENT SCIENTIST must sign both copies of this letter and return one signed copy to the PROVIDER. The PROVIDER will then send the MATERIAL.

**PROVIDER INFORMATION and AUTHORIZED SIGNATURE**

Provider Scientist: Arnold Kirshenbaum, M.D.
Dean Metcalfe, M.D.

Name of Authorized Official: Richard K. Williams, Ph.D.

Title of Authorized Official: Lead Technology Development Associate, Office of Technology Development, NIAID, NIH

Provider Organization: NIAID, NIH, HHS

Address for Notices: Office of Technology Development
6610 Rockledge Drive, Suite 2800
Bethesda, MD 20892 *(Zip Code for Courier: 20817)*

Certification of Authorized Official: This Simple Letter Agreementhas been modified. Modifications are attached.

Signature of Provider Authorized Official

Date: March 26, 2012

**RECIPIENT INFORMATION and AUTHORIZED SIGNATURE**

Recipient Scientist: Li Li, M.D.

Name of Authorized Official: Yanhong Zhu,MPA.

Title of Authorized Official: Science and education Dept.
Shanghai First People's Hospital

Recipient Organization: Shanghai First People's Hospital

Address for Notices: Dept. of Laboratory Medicine
85 Wujin Rd.
Shanghai, 200080

Signature of Recipient Authorized Officer

Date: 3/20/2012

Certification of Recipient Scientist: I have read and understood the conditions outlined in this Agreement and I agree to abide by them in the receipt and use of the MATERIAL.

Recipient Scientist

Date: 3/20/2012
